# Modelling human zygotic genome activation in 8C-like cells *in vitro*

**DOI:** 10.1101/2021.10.28.466259

**Authors:** Jasmin Taubenschmid-Stowers, Maria Rostovskaya, Fátima Santos, Sebastian Ljung, Ricard Argelaguet, Felix Krueger, Jennifer Nichols, Wolf Reik

**Affiliations:** Epigenetics Programme, Babraham Institute, Cambridge, UK; Centre for Trophoblast Research, University of Cambridge, Cambridge, UK; Wellcome-Medical Research Council Cambridge Stem Cell Institute, University of Cambridge, Cambridge, UK; Department of Physiology, Development and Neuroscience, University of Cambridge, Cambridge, UK; Wellcome Trust Sanger Institute, Hinxton, Cambridge, UK

## Abstract

The remodelling of the epigenome and transcriptome of the fertilised oocyte to establish totipotency in the zygote and developing embryo is one of the most critical processes in mammalian embryogenesis. Zygotic or embryonic genome activation (ZGA, EGA) in the 2-cell embryo in mouse, and the 8-cell embryo in humans, constitutes the first major wave of transcription. Failure to initiate ZGA leads to developmental defects, and contributes to the high attrition rates of human pre-implantation embryos. Due to limitations in cell numbers and experimental tractability, the mechanisms that regulate human embryonic genome activation in the totipotent embryo remain poorly understood. Here we report the discovery of human 8-cell like cells (8CLCs) specifically among naïve embryonic stem cells, but not primed pluripotent cells. 8CLCs express ZGA marker genes such as *ZSCAN4, LEUTX* and *DUXA* and their transcriptome closely resembles that of the 8-cell human embryo. 8-cell like cells reactivate 8-cell stage specific transposable elements such as *HERVL* and *MLT2A1* and are characterized by upregulation of the DNA methylation regulator *DPPA3*. 8CLCs show reduced SOX2 protein, and can be identified based on expression of the novel ZGA-associated protein markers TPRX1 and H3.Y *in vitro*. Overexpression of the transcription factor *DUX4*. as well as spliceosome inhibition increase ZGA-like transcription and enhance TPRX1+ 8CLCs formation. Excitingly, the *in vitro* identified 8CLC marker proteins TPRX1 and H3.Y are also expressed in 8-cell human embryos at the time of genome activation and may thus be relevant *in vivo*. The discovery of 8CLCs provides a unique opportunity to model and manipulate human ZGA-like transcriptional programs *in vitro*, and might provide critical functional insights into one of the earliest events in human embryogenesis *in vivo*.

**Highlights:**

- ZGA markers and transposable elements are expressed in 8CLCs among naïve human stem cells
- The transcription factor *DUX4* and spliceosome inhibition induce ZGA-like transcription
- 8CLC marker proteins TPRX1 and H3.Y are expressed in nuclei of 8-cell human embryos
- 8CLCs serve as a novel *in vitro* model for human ZGA

## Introduction

Mammalian embryogenesis begins shortly after fertilisation with the formation of the totipotent zygote. Totipotency is established through epigenetic and transcriptional remodelling and licenses formation of all cell types of the developing organism (Tarkowski, 1959). Zygotic or embryonic genome activation (ZGA, EGA) marks the first major wave of transcription in the totipotent mouse 2-cell and human 8-cell embryo, and is essential for the ensuing first lineage decisions (Aoki et al., 1997; Braude et al., 1988; Kigami et al., 2003; Latham and Schultz, 2001; Lee et al., 2014; Vassena et al., 2011). Failure to accurately remodel the epigenome or activate embryonic transcription contributes to substantial lethality of human pre-implantation embryos, but may also have longer term consequences later in development (Niakan et al., 2012). Understanding the molecular events regulating genome activation is therefore important for human reproduction and health. The direct study of human embryos, however, is practically and ethically limited.

To study mammalian early development *in vitro*, mouse and human pluripotent or embryonic stem cells (PSCs, ESCs) have been used as model systems (Evans and Kaufman, 1981; Thomson et al., 1998). Naïve human PSCs correspond to cells of the pre-implantation epiblast, whereas primed ones represent post-implantation stage cells (Huang et al., 2014; Nakamura et al., 2016; Theunissen et al., 2016). Moreover, among naïve mouse ESCs, a small subpopulation of so-called 2C-like cells (2CLCs) has been described, that closely resembles the totipotent 2-cell embryo *in vivo* (Macfarlan et al., 2012). Although these cells have already undergone ZGA, in an embryo context before their derivation from mouse blastocyst, they re-activate a 2-cell or ZGA-like transcriptional and epigenetic programme in culture, and have been shown to cycle in and out of this state from mouse ESCs (Macfarlan et al., 2012; Rodriguez-Terrones et al., 2018). 2CLCs can be identified based on the upregulation of the endogenous retrovirus MERVL, and can be used to study ZGA-like transcription *in vitro* (Alda-Catalinas et al., 2020; Eckersley-Maslin et al., 2016; Ishiuchi et al., 2015; Macfarlan et al., 2012). Moreover, 2CLCs have been reported to possess greater developmental potential than conventional ESCs, and are able to contribute to extra-embryonic lineages during mouse development (Macfarlan et al., 2012; Shen et al., 2021). To date no equivalent *in vitro* human 2 cell- or 8 cell-like cell type has been described.

Here we report the discovery of human 8-cell like cells, 8CLCs, specifically among human naïve ESCs. 8CLCs express ZGA marker genes such as *ZSCAN4, LEUTX, TPRX1* and *PRAMEFs*. Their transcriptome, including transposon expression profile, closely resembles that of human 8-cell embryos. 8CLCs are characterized by upregulation of the DNA methylation regulator *DPPA3*, and lower SOX2 protein levels. 8CLCs can be identified *in vitro* by the expression of the novel protein marker TPRX1. Overexpression of the transcription factor *DUX4* not only increases ZGA-like gene expression, but also enhances TPRX1-positive 8CLCs formation. We have thus uncovered a novel *in vitro* cell state and model system, 8CLCs, that allows us to study and manipulate human ZGA-like transcription *in vitro*.

## Results

### ZGA markers are expressed in 8CLCs among naïve human PSCs

To study functionally the molecular events of human ZGA regulation we sought to discover an *in vitro* model system. We defined ZGA markers from human embryo RNA sequencing data as well as from overexpression studies of the ZGA-associated transcription factor (TF) DUX4 in different cell lines (De Iaco et al., 2017; Hendrickson et al., 2017; Petropoulos et al., 2016) (Figure 1A). From human embryo transcriptome data, we chose features that were overrepresented at the 8-cell stage during genome activation (embryonic day 3, E3) (Petropoulos et al., 2016) (Figure S1A). These features include known ZGA markers such as the transcription factor *ZSCAN4* (Falco et al., 2007; Zalzman et al., 2010), the eutherian specific paired-like homeodomain protein *LEUTX* (Jouhilahti et al., 2016), as well as the *TRIM* and *PRAMEF* superfamily of genes, all of which are specifically expressed during the 8-cell stage (Figure S1A-S1D). To refine this gene set we integrated *DUX4* overexpression data. DUX4 is one of the few transcription factors that has been implicated in mouse and human ZGA (De Iaco et al., 2017; Hendrickson et al., 2017). It is specifically expressed in the 4-cell human embryo, just prior to ZGA initiation, and has been suggested to act as an activator of ZGA-like transcription. Endogenous upregulation as well as overexpression of exogenous *DUX4* in different human primary cells and cell lines leads to the upregulation of a common set of genes, almost all of which are expressed specifically at the time of human ZGA (Figures S1E and 1F) (Hendrickson et al., 2017; Jiang et al., 2020; Yao et al., 2014). We combined the human embryo *in vivo* data with the *DUX4* overexpression studies to define a common set of human ZGA markers (Table S1) that can be studied *in vitro*.

**Figure 1.**
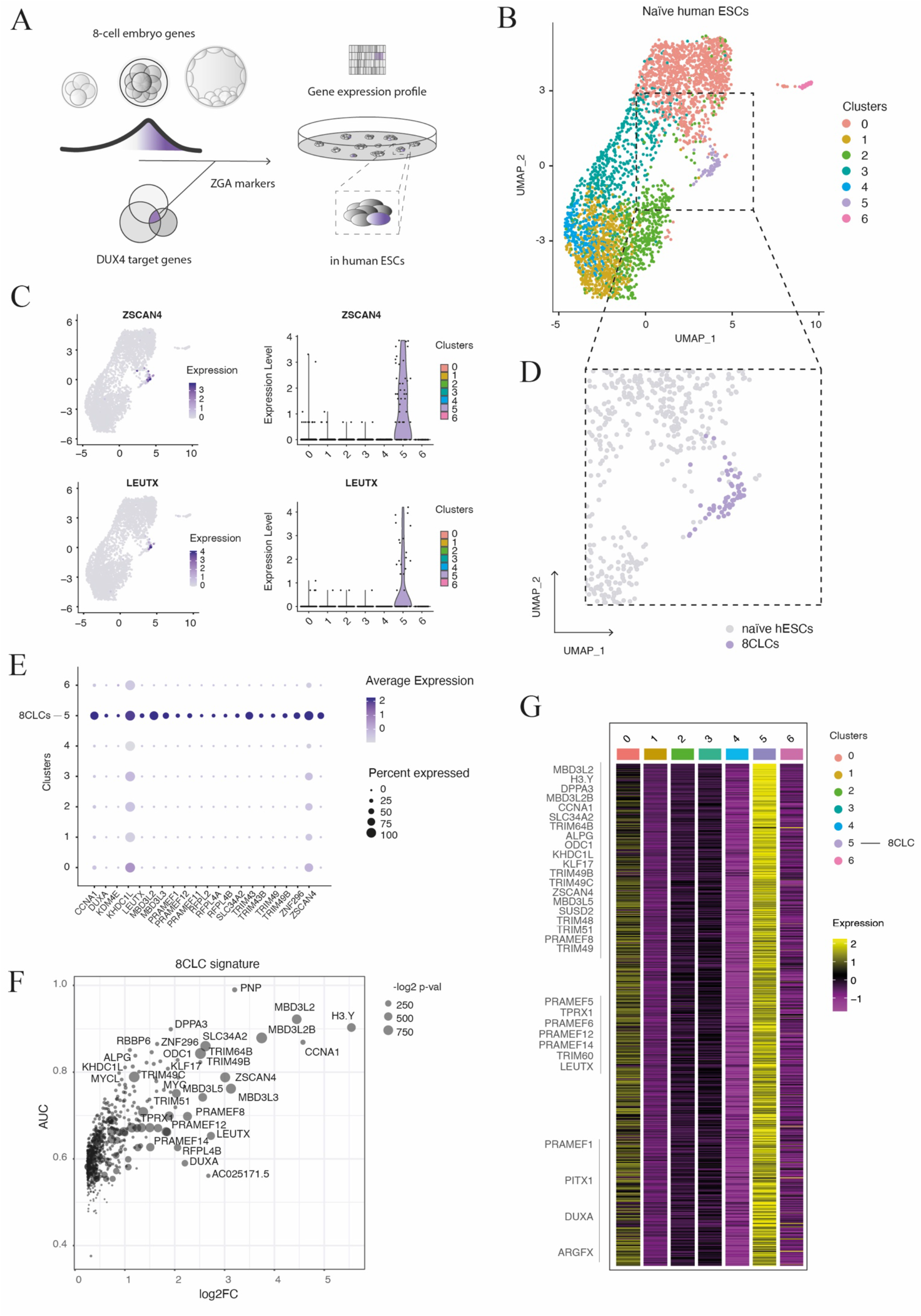
ZGA marker expression in 8CLCs and naïve hESCs. (A) Schematic of establishing a novel *in vitro* model system for human ZGA. (B) UMAP of naïve HNES1 hESCs cultured in PXGL. Cell clustering is based on normalized, scaled single cell RNA expression data. (C) Normalized scaled gene expression of *ZSCAN4* and *LEUTX* in naïve human ESCs, visualized using UMAPs (left) or Violin plots of clustered cells (right) (clusters 0, 1, 2, 3, 4, 5, 6 contain n = 1237, 709, 611, 517, 251, 55, 30 cells respectively). (D) UMAP of cluster 5 cells, highlighted as ‘8C-cell like cells’, ‘8CLCs’, comprising of ~1.6% (55 of 3410) of total naïve cells. (E) Dotplots of ZGA marker frequency and average expression level of clustered naïve hESCs. (F) Gene expression signature of cluster 5 cells, or 8CLCs, as determined by the ‘Findmarkers’ function in Seurat. AUC, area under curve; LOG2FC, log2 fold-change; -log2 p-val, negative log2 p-value. (G) Heatmap of normalized, scaled average expression of 8CLC (cluster 5) markers (rows) in naïve hESC clusters (columns, cluster 0 – 6).

We next assessed the presence of these markers in single-cell transcriptome data of cultured human embryonic stem cells. We first assessed ZGA-like transcription in human embryo-derived naïve HNES1 cells cultured under PXGL conditions (Guo et al., 2017; Guo et al., 2016; Rostovskaya et al., 2019). Single-cell expression data revealed several distinct populations of cells as judged by dimensionality reduction and clustering (Figure 1B). Notably, a distinct subset of 55 cells out of 3410 (cluster 5 in our dataset) showed clear upregulation of several ZGA-like transcripts simultaneously (Figures 1C and 1D). These transcripts did not only include ZGA genes such as *LEUTX* and *ZSCAN4*, but also extended to most previously identified ZGA markers, including *DUXA, MBD3L3*, and *TRIM49* (Figure S2A, S2B). The level of ZGA gene expression and percentage of cells expressing those markers varied within the cluster (Figures 1D and 1E), but was in general consistently high in cells belonging to cluster 5 (Figures S2C – S2F). We termed cluster 5 cells ‘8-cell like cells’, or ‘8CLCs’.

8CLCs can be discriminated from the remaining naïve ESCs based on the expression of more than 700 markers that are specifically upregulated in cluster 5 (Figures 1F and 1G). This 8CLC signature includes all previously selected ZGA markers *(ZSCAN4, LEUTX, PRAMEF1, MBD3L3* etc) but notably also additional factors such as the maternal and zygotic DNA demethylation regulator *DPPA3* (Huang et al., 2017), the naïve pluripotency marker *KLF17* (Blakeley et al., 2015; Guo et al., 2016), the histone variant *H3.Y* (a recently described *DUX4* target) (Resnick et al., 2019), and the eutherian specific genome activation associated factor *TPRX1* (Madissoon et al., 2016; Maeso et al., 2016) (Figure 1F, Table S2). 8CLC signature genes are not detected or expressed at low levels in the other subpopulations of clustered naïve ESCs (Figure 1G). These analyses show that 8CLCs are characterised by a distinct and unique gene expression signature within a subpopulation of naïve human pluripotent stem cells.

We next asked if the presence of 8CLCs was specific to the embryo-derived HNES1 cell line (under PXGL culture conditions) or if they could be found in other naïve hPSCs as well. We analysed single cell expression data of human H9-derived NK2 ESCs that were genetically reprogrammed from primed cells via overexpression of *NANOG* and *KLF2* and cultured in t2iLGö (Messmer et al., 2019). Indeed, naïve NK2 cells also contained a distinct subpopulation of 8CLCs that upregulate ZGA markers *(TRIM49, MBD3L3 and LEUTX)* and 8CLCs signature genes *(DPPA3* and *SUSD2)* as compared to the remaining naïve hPSCs (Figures S2G and S2H). In both single cell RNA-sequencing datasets ZGA-like transcriptome expressing cells comprise around 1.5% of the population of total naïve cells, a similar proportion to mouse 2C-like cells (Macfarlan et al., 2012).

We further compared ZGA-like transcription under different naïve and primed culture conditions (Theunissen et al., 2016). In addition to PXGL cultured cells, 5iLA, 4iLA and t2iLGö cells are considered truly naïve as they resemble cells of the human pre-implantation epiblast *in vivo*. NHSM media, primed culture conditions (WIBR cells), and differentiated neuronal precursor cells (NPCs) represent later stages in development (Theunissen et al., 2016) (Nakamura et al., 2016). ZGA markers and most 8CLC signature genes are found highly expressed in naïve 5iLA, 4iLA, and t2iLGö cells, but are lower in pseudo-naïve (NHSM), primed (WIBR) and differentiated cells (NPCs) (Figure S3A). Similarly, ZGA markers and 8CLCs signature are downregulated from naïve (PXGL) to primed (E8 or XAF – containing XAV939, Activin A and FGF2, similar to mouse EpiSC medium) transition of HNES1 cells, as well as chemically reset cR-H9-EOS cells (Figure S3B) (Rostovskaya et al., 2019) (Sumi et al., 2013). This data suggests that 8CLCs and ZGA-like transcription can be found in naïve state PSCs but not primed or differentiated cells.

### 8CLC transcription resembles that of human 8-cell embryos

We next compared the 8CLC transcriptome *in vitro* to human 8-cell embryos *in vivo*. We analysed gene expression profiles at different stages and from different studies of human pre-implantation development (Petropoulos et al., 2016; Stirparo et al., 2018; Xue et al., 2013; Yan et al., 2013). These analyses showed that both ZGA markers and 8CLC signature genes peak in 8-cell human embryos during ZGA (E3 embryos) and are downregulated thereafter in morulae (E4 embryos) (Figures 2A, 2B, S3C and S3D). By contrast, while the top 8CLC signature genes are upregulated during ZGA (Figure S4A), naïve marker genes that are lowly expressed in 8CLCs are downregulated during ZGA and are only expressed at later stages in blastocysts (E5 – E7) (Figure S4B). Moreover, comparison of ZGA markers in single cell expression data of conventional, naïve, 8CLCs and the human embryo, shows strong similarity between 8CLCs and the 8-cell stage embryo (E3) (Figure 2C), with almost half of 8CLC signature genes overlapping with 8-cell human embryo markers (Figure 2D). Finally, when we combined single cell sequencing data of naïve ESCs and human embryos, most 8CLCs clustered together with 8-cell stage embryo (E3) and morula cells (E4), while naïve hESCs clustered together with blastocyst stage cells (E5 – E7) (Figures 2E and 2F). These analyses confirm a remarkable similarity in gene expression patterns of 8CLCs *in vitro* and human 8-cell embryo *in vivo*.

**Figure 2.**
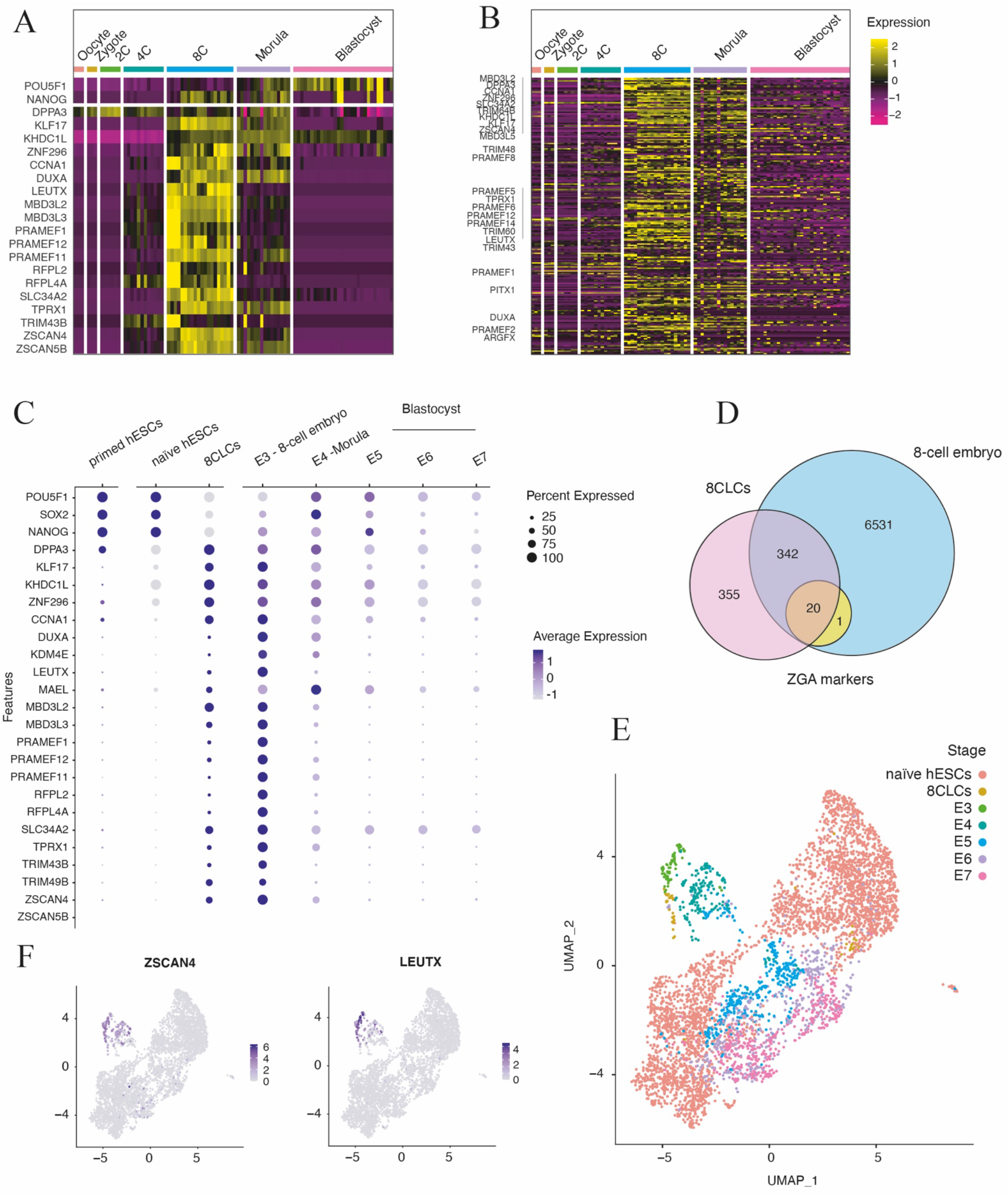
8CLCs transcriptome signature in human embryos. (A) Heatmap of normalized scaled expression of ZGA markers and pluripotency genes, as well as (B) 8CLCs signature genes (rows) in human pre-implantation embryo cells (columns) (Yan et al., 2013). (C) Dotplots of frequency and average expression of ZGA markers and pluripotency genes in 8CLCs, naïve hPSCs, primed hPSCs (Rostovskaya et al., 2019) and human 8-cell (E3) to blastocyst (E7) stage embryos (Petropoulos et al., 2016). (D) Overview of shared ZGA markers (n=21), 8CLCs genes (n=717) and 8-cell embryo markers (n=6894) (Petropoulos et al., 2016). (E) Clustering of individual 8CLCs, naïve hPSCs and 8-cell to blastocyst stage embryo cells (E3 – E7) (Petropoulos et al., 2016) depicted on a UMAP. Datasets have been combined and merged in Seurat. (F) *ZSCAN4* and *LEUTX* expression levels in clustered 8CLCs, naïve hESCs and human embryos (E3 – E7) (Petropoulos et al., 2016).

### ZGA-specific TF motifs and transposable elements are enriched in 8CLCs

We next wanted to identify candidate TF signatures that might regulate ZGA-like transcription. We analysed potential binding sites in the genome surrounding (+/− 10kb) 8CLC signature genes and identified several enriched motifs, such as *DUX4, DUXA, KLF17* (Figures 3A and 3B). Notably, *KLF17* is upregulated in 8CLCs (Figure 3C) and 8-cell human embryos (Figure 2B) (Blakeley et al., 2015). *DUX4, DUXA, KLF17* binding motifs have also been reported to be highly accessible during human ZGA (Bentsen et al., 2020; Liu et al., 2019). Thus, transcriptional regulation of ZGA signature genes *in vivo* could potentially also regulate 8CLC-formation *in vitro*.

**Figure 3.**
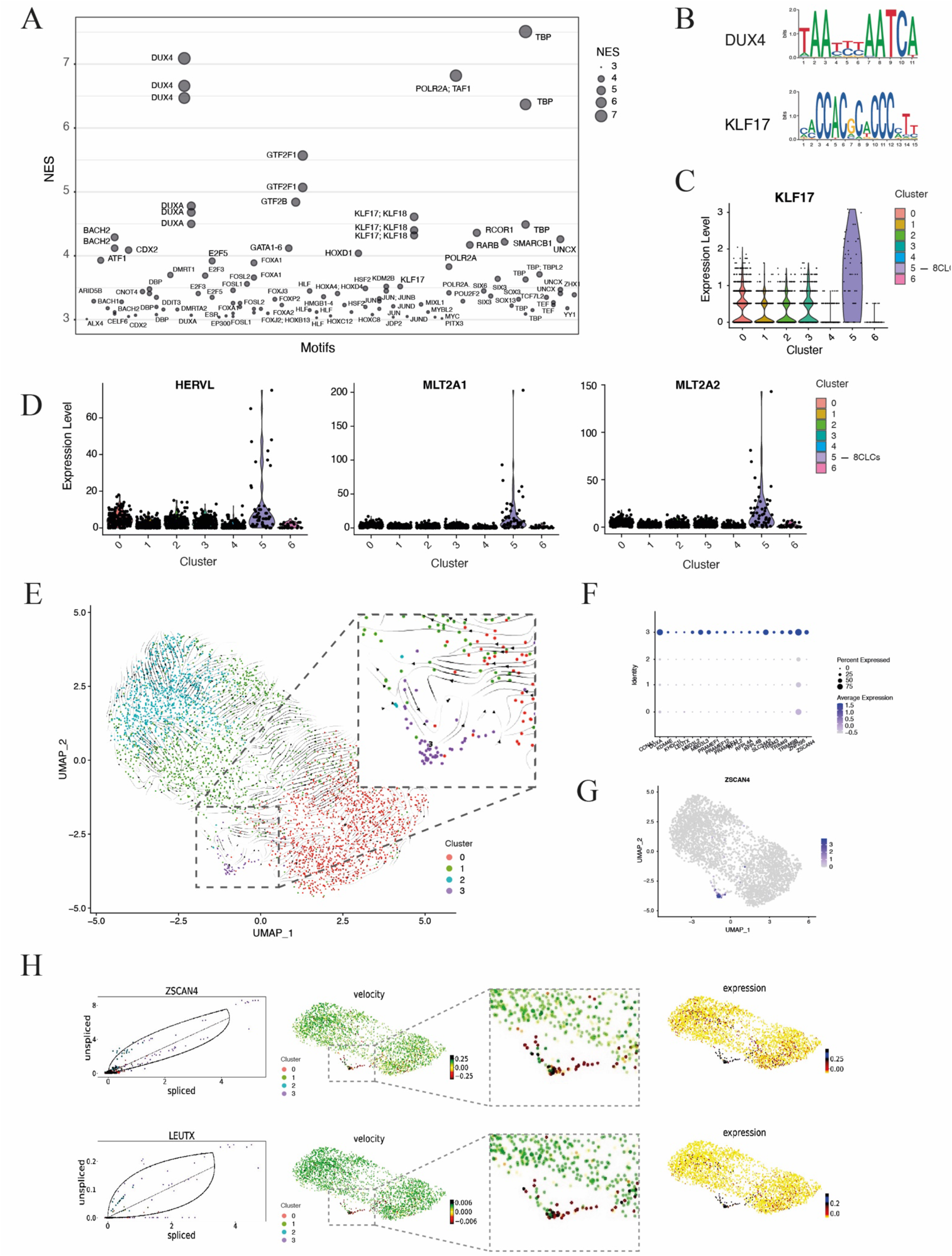
TF motifs and transposon expression in 8CLCs. (A) TF motifs identified around transcriptional start sites (TSS ± 10kb) of the top 200 8CLCs signature genes using standard parameters in RcisTarget. NES; Normalized Enrichment Score. (B) DUX4 and KLF17 binding motifs as identified in (A) are shown. (C) Violin plots of normalized, scaled *KLF17* expression in clustered naïve human ESCs. (D) Raw reads of transposable elements in clustered naïve ESCs derived from single cell RNA-seq data. (E) RNA velocity in naïve hESCs and 8CLCs; cluster 3 represents 8CLCs – see also (F) ZGA marker and (G) *ZSCAN4* expression in clustered hESCs and 8CLCs. (H) RNA velocity analysis of ZGA markers, such as *ZSCAN4* and *LEUTX*, in 8CLCs and naïve hESCs. Left panel: steady-state ratio (black diagonal line), overall dynamics (black curve) and ratio of unspliced vs. spliced mRNA in single cells, coloured according to their cluster identity; middle panel: RNA velocity of marker genes; right panel: expression levels of markers.

We also assessed the transposon expression landscape in 8CLCs. Endogenous retroviral or transposable elements such as MERVL have been described to regulate mouse 2-cell embryo development and 2C-like cell transcription *in vivo* and *in vitro* (Kigami et al., 2003; Peaston et al., 2004; Percharde et al., 2018; Svoboda et al., 2004). While human LINEs, SINEs and DNA transposons were detected at similar levels in both 8CLCs and naïve hESCs by bulk analysis, there was a trend for higher LTR element expression in 8CLCs (Figure S4C). We thus further analysed LTR sub-families and found slightly higher levels of *LTR-ERVL* and *LTR-ERVK*, but not other LTRs in 8C-like cells (Figure S4D). Remarkably, specific upregulation of *HERVL, MLT2A1* and *MLT2A2* transcripts was detectable in some 8CLCs (cluster 5) but not naïve hESCs (Figures 3D). *MLT2A1, MLT2A2* and *HERVL* are the most strongly upregulated repeats in the 8-cell human embryo *in vivo* and gain chromatin accessibility specifically during human ZGA (Liu et al., 2019). This shows that conserved ZGA-specific transposable elements are activated in some 8CLCs *in vitro*, similar to their upregulation in the embryo *in vivo*.

We next explored the gene expression dynamics in naïve hESCs and 8CLCs. We used single cell RNA velocity to compare immature to mature mRNA species and predict future RNA expression patterns and direction of cell fate transitions (Figure 3E). These analyses suggest bi-directional transitions from naïve ESCs to 8CLCs and from 8CLCs towards the naïve state (see arrows indicating direction of RNA velocity in cluster 3 cells, Figure 3F – 3E). This bi-directionality was also detectable in ZGA markers, such as *ZSCAN4* and *LEUTX*, which displayed an increase in RNA velocity in 8CLCs (higher unspliced to spliced ratio), corresponding to their upregulation, as well as a decrease, so downregulation, from 8CLCs towards the naïve hESCs state (Figure 3H and S4E). These data suggest dynamic gene expression changes and transitions of 8CLCs to and from naïve hESCs, pointing towards a cycling nature of 8CLCs, similar to mouse 2CLCs.

### 8CLCs are marked by TPRX1 protein expression

We next wanted to define protein markers that can be used to identify 8CLCs *in vitro*. While many genes are upregulated in 8CLCs (Figure S5A), we identified specifically one protein that is highly expressed in 8CLCs in culture, TPRX1 (Figure 4A). *TPRX1* (tetrapeptide repeat homeobox 1) is a protein coding gene of the paired (PRD)-like homeobox gene family of transcription factors that have been previously implicated in ZGA-like transcription; its biological role, however, is unknown (Madissoon et al., 2016; Maeso et al., 2016). While TPRX1 positive cells were detected among naïve HNES1 cells in culture, no primed H9 PSCs stained positive for the marker and EpiLCs also appear negative for TPRX1 (Figure 4A, S5B). TPRX1 expressing cells were found among reprogrammed naïve H9 PSCs (WA09-NK2), and human fibroblast-derived naïve iPSCs (FiPSCs) of non-embryonic origin, both grown under t2iLGö conditions (Takashima et al., 2014), as well as HNES1 cells grown in 4iLA and 5iLA (Theunissen et al., 2014) (Theunissen et al., 2016) (Figure S5C). Analysis of stemness markers revealed that TPRX1-positive 8CLCs show reduced levels of the pluripotency marker SOX2, both on the transcriptional (Figure S5D) as well as protein level (Figures 4B, S5E and S5F). We further analysed DNA methylation levels and found that TPRX1+ cells have similar methyl CpG (5mCpG), but increased 5-hydroxymethyl cytosine (5hmC) levels as compared to naïve hESCs (Figure S5G – S5I). Moreover, some TPRX1 positive cells also display high levels of the DUX4 target gene H3.X/Y, both at the single cell RNA (Figure S5A), as well as at the protein level via immunofluorescence (Figure 4C). These results show that TRPX1, together with the histone variant protein H3.X/Y, can be used to identify 8CLCs among naïve human PSCs in culture.

**Figure 4.**
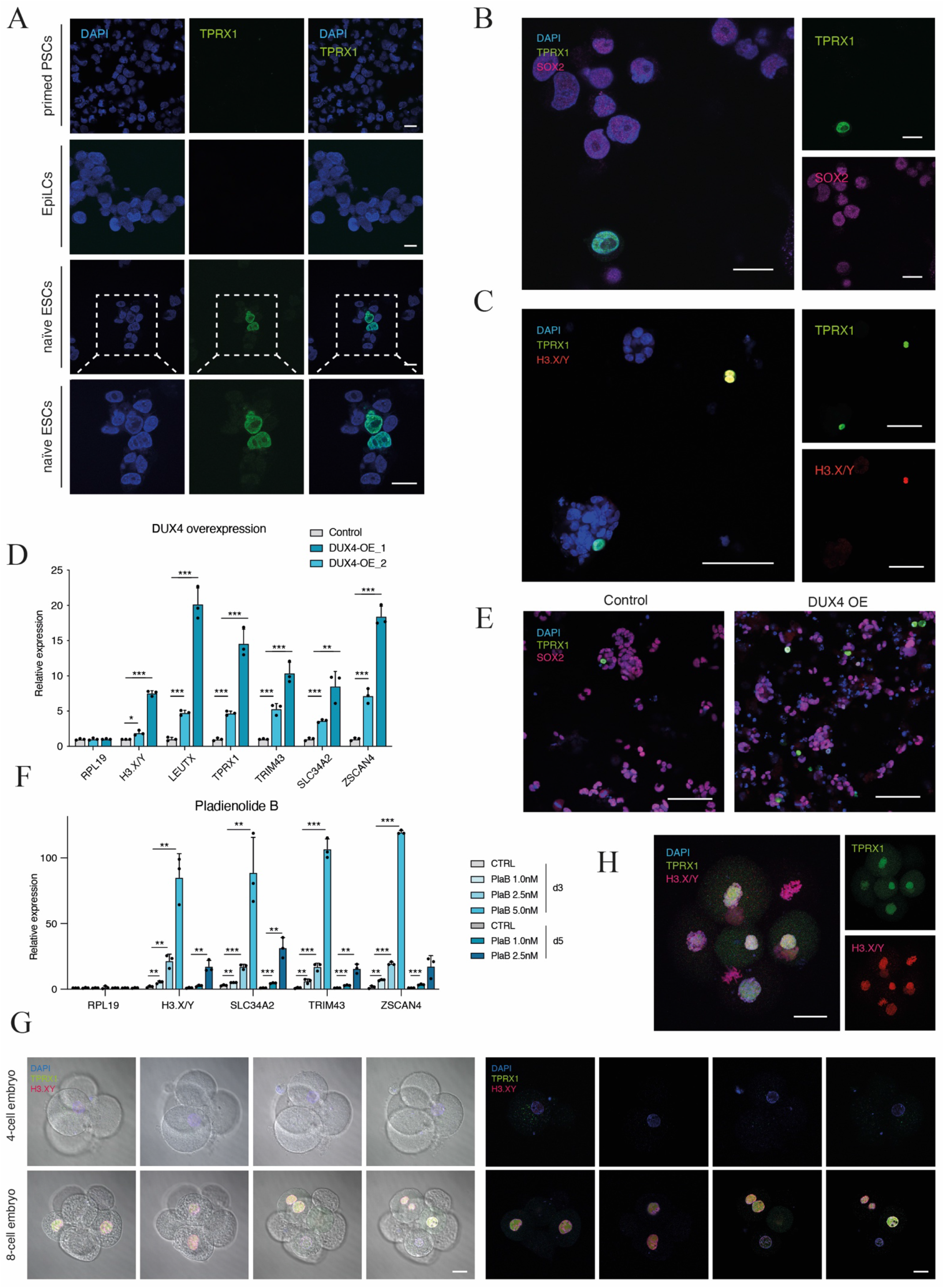
8CLCs are marked by TPRX1 protein expression. (A) Immunofluorescence (IF) staining of cytospun naïve ESCs, EpiLCs and primed hPSCs for TPRX1. Scale bar, 15μm. (B) IF staining of TRPX1 and SOX2 in cytospun naïve HNES1 cultured in PXGL. Scale bar, 20μm (C) Representative images of H3.X/Y and TPRX1 stained HNES1 ESCs plated onto coverslips. Scale bar, 100μm. (D) Expression levels of ZGA marker genes assessed by RT-qPCR in two biological replicates of *DUX4* overexpressing HNES1 cells (*DUX4* OE) as compared to control cells (WT). Data are shown as mean ± SD (n = 3 technical replicates) of fold-change compared to control HNES1, and are representative of three independent experiments. p-value *P<0.05, **P<0.01, ***P<0.001, absence of stars (non-significant, ns): p-value>0.05; unpaired, two tailed Student’s t-test. (E) IF staining of TPRX1 and SOX2 in *DUX4* overexpressing naïve HNES1 and control cells in PXGL plated onto coverslips. Scale bar, 100μm. (F) ZGA marker gene expression measured by RT-qPCR in Pladienolide B (PlaB 1.0-5.0nM, 3-5 days) and vehicle treated control cells. Data are shown as mean ± SD (n = 3 technical replicates) of fold-change compared to control, and are representative of three independent experiments. p-value *P<0.05, **P<0.01, ***P<0.001, absence of stars (non-significant, ns): p-value>0.05; unpaired, two tailed Student’s t-test. (G) Brightfield overlays and immunofluorescence images of TPRX1- and H3.X/Y-stained human pre-implantation embryos at the 4-cell and 8-cell stage. Scale bar, 25μm. (H) Immunofluorescence image of dividing human 8-cell stage blastomeres stained for TPRX1 and H3.X/Y. Scale bar, 25μm.

Next, we wanted to see if 8CLC formation can be modified in culture. Since our motif analysis identified DUX4 binding sites enriched in 8CLC signature genes, we asked if *DUX4* overexpression would alter 8CLCs abundance, and hence introduced a codon-adapted version of the TF into naïve hESCs (Jagannathan et al., 2016). Overexpression of *DUX4* did not only substantially increase transcription of ZGA marker genes, such as *LEUTX, ZSCAN4, TPRX1* and *TRIM43* (Figure 4D), but also up to 20-fold increased the number of TPRX1 positive, lower SOX2 expressing 8CLCs among naïve hESCs in culture (Figures 4E and S6A – S6C). Similar to wildtype 8CLCs, TPRX1+ cells generated via *DUX4* overexpression also harbour higher 5hmC levels (Figure S6D and S6E), some of which were also positive for H3.X/Y (Figure S6F) and notably, were actively dividing (Figure S6G). Additionally, induction of *DUX4* and subsequent withdrawal of transgene expression led to consequential downregulation of ZGA genes after the initial upregulation (Figure S6H), thus indicating that ZGA-like transcription is likely reversible and transient in naïve hESCs. We characterized the exogenous *DUX4* (*DUX4-CA*) expressing cells in more detail transcriptionally and epigenetically. While we could not reliably detect endogenous *DUX4* due to its very low expression levels, we identified *DUX4-CA* as a novel marker of 8CLCs among naïve hESCs in our data (Figure S6I – S6K). Using single cell multi-omics (combined scRNA-seq and scATAC-seq) we also sorted these cells computationally (based on their transcriptional signature) and have thus been able to generate chromatin accessibility profiles of 8CLCs, where we observed increased accessibility at sites proximal to ZGA-markers, such as *ZSCAN4* and *RFPL4A*, as compared to naïve hESCs (Figure S6L). This increased accessibility is also observed in 4- or 8-cell embryos prior and around the time of genome activation (Figure S6L). In addition, we tested a recently reported spliceosome inhibitor, Pladienolide B, that has been shown to increase ZGA-like transcription and developmental potency in mouse stem cells (Shen et al., 2021). We observed a strong, dosage-dependant upregulation of ZGA genes such as *ZSCAN4* and *H3. Y* in human ESCs, pointing towards a conserved role of spliceosome regulation in totipotency (Figure 4F). These results highlight that both ZGA-like transcription and *in vitro* 8CLC-formation can be increased through genetic interference, such as *DUX4* overexpression, as well as pharmacological inhibition, such as Pladienolide B treatment and can thus be used to study ZGA-like properties *in vitro*.

Finally, we tested if the newly identified 8CLC markers TPRX1 and H3.Y can also be detected in human pre-implantation stage embryos. Remarkably, 8-cell human embryos stained strongly positive for TPRX1 and H3.Y at the time of human embryonic genome activation, but these markers were not detectable in 4-cell human embryos (Figure 4G). Interestingly, while TPRX1 was not present in blastomeres undergoing cell division, reassuringly the histone variant H3.Y showed strong association with prometa- and metaphase chromosomes in dividing cells (Figure 4H). These findings confirm the *in vitro* identified 8CLCs specific proteins TPRX1 and H3.Y as novel markers of genome activation in human 8-cell embryos.

## Discussion

To analyse and manipulate the molecular events underlying genome activation, an *in vitro* model of human ZGA is needed. Such a model would not only have practical implications for our ability to study pre-implantation development functionally, but would also improve our knowledge of human reproduction and health.

In this study we report that human ZGA-like transcription, which occurs in the 8-cell embryo *in vivo*, can be found in a distinct population, approximately 1.6% of naïve stem cells, *in vitro*. We termed these cells ‘8-cell like cells’ (8CLCs). 8CLCs express ZGA markers such as *ZSCAN4, LEUTX* and *PRAMEFs*, and are characterized by an 8CLC specific transcriptome signature. This signature shows increased expression of novel factors such as *DPPA3, H3.Y* and *KLF17*, and distinguishes 8CLCs from naïve PSCs. Our discovery of human 8CLCs as the first ZGA-stage cell population in a species other than mouse, with a comparable frequency (1-5% 2CLCs among mouse ESCs), suggests that this rare transcription event may be an intrinsic property of pluripotent stem cells, rather than a randomly emerging one. The activation of transposable elements in 8CLCs strikingly similar to human 8-cell embryos also points towards conserved transcriptional networks in stem cells and embryos – akin to other repeat element driven cell states that have been reported (Grow et al., 2015; Pontis et al., 2019; Wang et al., 2014). As 8CLCs specifically activate transposable elements such as *MLT2A1* and *MLT2A2*, which both harbour DUX4 binding sites, it will be important to assess their functional role in 8CLC formation and ZGA-like transcription *in vitro*.

We further identified TPRX1 as the first 8CLC marker protein that allows monitoring of ZGA-like transcription in cells *in vitro* as well as in 8-cell human embryos. *TPRX1* is itself an ETCHbox gene (eutherian-specific homeobox genes, which also include *LEUTX, ARGFX* and *DPRX)* and has been reported to be expressed in human pre-implantation embryos transcriptionally (Jouhilahti et al., 2016; Maeso et al., 2016). Although *TPRX1* has been implicated in ZGA-like transcription before, its general biological function as well as specifically during ZGA, still needs to be elucidated.

Similarly, the second 8CLC marker that we described, H3.Y (Resnick et al., 2019), which is also expressed in the 8-human cell embryo, and strongly upregulated upon *DUX4* induction *in vitro*, is a good candidate to study chromatin remodelling events involved in ZGA-like transcription. Importantly, the discovery that both *in vitro* identified 8CLC marker proteins TRPX1 and H3.Y are expressed in nuclei of ZGA-stage 8-cell human embryos highlights their potential *in vivo* relevance and encourages further studies.

Interestingly, while 8CLCs upregulate some naïve pluripotency markers such as *DPPA3* and *KLF17*, they downregulate other stem cell markers including SOX2, also at the protein level. It will be important to see if SOX2 downregulation occurs only transiently and is a cause or consequence of 8CLC formation, and how compatible this reduction is with 8CLC survival and growth as compared to the pluripotency state in naïve ESCs and how these changes in transcription are reflected in cellular properties, such as differentiation potential. Moreover, it remains to be assessed whether 8CLCs possess increased developmental competence, similar to mouse 2CLCs, and would more readily contribute to extraembryonic tissues in an embryo environment.

Altogether, we have discovered a novel cell state which enables the functional and molecular characterisation of ZGA-like transcription *in vitro*. The appearance, maintenance and enrichment of 8CLCs will allow the study of genome activation in culture and make it amenable to genetic and pharmacological manipulation in a high-throughput way. Discoveries made from studying ZGA-like transcription *in vitro* may also provide significant insights into the regulation of genome activation during human preimplantation development and may have vital consequences for reproduction, health and technology.

### Limitations

Although 8CLCs will be useful to study human genome activation in a non-invasive, accessible and systematic way *in vitro*, the validation of findings in this model system still depends on work in human embryos. Furthermore, while ZGA-like marker expression in 8CLCs and 8-cell blastomeres is highly similar, these two cell states expectedly differ as well. This is most likely due to differences in origin and maintenance (i.e. *in vitro* passaged cell lines as compared to directly fertilization-derived embryos) as well as developmental potential (the ability to differentiate in tissue culture as compared to the capacity to generate a fully grown organism) of the two cell types. Moreover, while the genome is originally activated in human 8-cell embryos, 8CLCs have already undergone ZGA and are re-establishing ZGA-like programs *in vitro*.

While most clustered 8CLCs are characterized by a defined transcriptional signature, certain markers are expressed to varying degrees in individual cells (Figure 1E). *ZSCAN4*, for example, is upregulated in most 8CLCs (see Figure S2A, S2B), while other markers, such as *TPRX1, DUXA* or the transposable elements, such as *MLT2A1*, are less broadly expressed (Figure S5A, Figure 3D). Similarly, partly overlapping markers, such as *Mervl* and *Zscan4*, have also been previously reported in mouse 2CLCs. Exploring the heterogeneity of 8CLCs will be important to further understand intermediate states and the cellular mechanism underlying the formation of 8CLCs in culture.

Relatedly, the stability of 8CLCs requires further assessment and comparison to the mouse system, where 2CLCs have been reported to transiently cycle into and out of a ZGA-like state. While we did not observe any changes in 8CLC percentage over time (>25 passages), we noticed that the upregulation of ZGA markers upon *DUX4* induction was only transient and reversible upon withdrawal. These results, together with the dynamic properties of 8CLCs in the RNA velocity analysis, point towards a cycling nature of 8CLCs in culture. More experimental evidence of the stability of 8CLCs (i.e., stable, metastable or transient) and their ability to revert back to the naïve state (as well as their conversion rate) will be required to answer such questions definitely.

To utilize 8CLCs to their full potential, some technical hurdles still need to be overcome. For example, to isolate this subpopulation of cells for downstream applications that are not single cell based, the identification of surface markers or the generation of endogenous reporter lines, similar to mouse 2CLC reporters, will be required. Also, enrichment of 8CLCs through adapted media compositions or altered culture conditions might allow increasing the numbers to obtain sufficient material for larger-scale experiments. Once isolation of 8CLCs can be done more easily, analyses of cell cycle and growth rates might help investigate origin and fate of 8CLCs in culture.

## Methods

### Human pluripotent stem cells lines

Human naïve WA09-NK2 H9 PSCs and naïve FiPSCs were kindly provided by Austin Smith (Takashima et al., 2014).

### Pluripotent stem cells culture

Naïve stem cells were cultured in a humidified incubator at 5% O2, 5% CO2 and 37°C. They were maintained on irradiated mouse embryonic fibroblast feeder cells (p2 DR4 expanded MEFs from WT-MRC Cambridge Stem Cell Institute) and were passaged every 2-3 days using TrypLE (Thermo Fisher Scientific, 12605028). ROCK inhibitor (10μM; Y-27632, 688000, Millipore) was added for 24h after passaging. Geltrex (hESC-qualified, Thermo Fisher Scientific, A1413302) was optionally added to the medium during re-plating on feeders (0.5μl per cm2 surface are), or without feeder cells (1.0μl per cm2).

Embryo-derived HNES1 naïve stem cells were cultured in PXGL culture conditions: N2B27 (1:1 DMEM/F12:Neurobasal, 0.5x N2 supplement, 0.5x B27 supplement, 1x nonessential amino acids, 2mM l-Glutamine, 1x Penicillin/Streptomycin (all from ThermoFisher Scientific), 0.1mM β-mercaptoethanol (Sigma-Aldrich), supplemented with 1μM PD0325901 (WT-MRC Cambridge Stem Cell Institute), 2μM Gö6983 (Tocris Bio-Techne, 2285), 2μM XAV939 (Tocris Bio-Techne, 3748) and 10ng/ mL human LIF (WT-MRC Cambridge Stem Cell Institute) (Guo et al., 2016), (Bredenkamp et al., 2019). For 5iLA media conditions, HNES1 were cultured in: N2B27 (see above), supplemented with 1μM PD0325901 (WT-MRC Cambridge Stem Cell Institute), 1μM IM-12 (Sigma-Aldrich), 0.5μM SB590885 (SigmaAldrich), 1μM WH-4-023 (A Chemtek), 10μM Y-27632 (Cell Guidance Systems), 50μg/ml bovine serum albumin (ThermoFisher Scientific), 0.5% KnockOut Serum Replacement (KSR; ThermoFisher Scientific), 20ng/ml Activin A (WT-MRC Cambridge Stem Cell Institute) and 10ng/mL human LIF (WT-MRC Cambridge Stem Cell Institute). 4iLA media was the same as 5iLA but with the omission of 1μM IM-12, the GSK3i (Theunissen et al., 2016; Theunissen et al., 2014). WA09-NK2 H9 and fibroblast derived FiPSCs naïve stem cells were maintained in t2iLGö: N2B27 (see above), supplemented with 1μM PD0325901 (WT-MRC Cambridge Stem Cell Institute), 2μM Gö6983 (Tocris Bio-Techne, 2285), 1μM CHIR99021 (WT-MRC Cambridge Stem Cell Institute) and 10ng/mL human LIF (WT-MRC Cambridge Stem Cell Institute) (Takashima et al., 2014). Primed WA09-NK2 PSCs were maintained in Essential 8 (E8) medium (Thermo Fisher Scientific A1517001) on Matrigel (Corning™ Matrigel™ Basement Membrane Matrix, Thermo Fisher Scientific 15575729).

### DUX4 overexpression in HNES1

A codon adapted version of human DUX4 (pCW57.1-DUX4-CA, Addgene plasmid #99281) was cloned into an inducible, BFP containing vector harbouring IR (inverted repeats) compatible with PBase (PiggyBac transposase). For inducible DUX4 expression, 1×10^6 HNES1 cells were plated onto Geltrex with ROCK inhibitor for 24h-48h, and electroporated using the NEON Transfection System and 100μl kit (Thermo Fisher Scientific, MPK10096) with 3ug PBase plasmid and 3ug vector. The conditions used for the NEON electroporation were the following: 1150V, 30ms pulse width, 2 pulses. Stable BFP expressing cells were sorted using FACS after >72h and expanded. DUX4 expression was induced by Doxycycline for 2-4h (2μg/ mL).

### Thawing, immunohistochemistry staining and imaging of human embryos

Cryopreserved 4-8 cell stage human embryos, left over from assisted conception programmes and kindly donated with informed consent to Human Fertilisation and Embryology Authority licence R0178, were thawed using EmbryoThaw (FertiPro) according to manufacturer’s instructions. After recovery in Cleave culture medium for an hour, zonae pellucidae were removed by brief incubation in acid Tyrodes solution and the embryos immediately fixed in 4% paraformaldehyde for 15 minutes. Immunofluorescence staining was performed as described previously (Roode et al., 2012). Briefly, embryos were incubated for 15 minutes in PBS containing 3 mg/mL polyvinylpyrrolidone (PBS/PVP, P0930, SigmaAldrich), transferred to PBS/PVP+ 0.25% Triton X-100 (23,472-9, SigmaAldrich) for permeabilization for 30 minutes. Blocking for at least 1 hour, and all subsequent procedures were performed using PBS containing 0.1% BSA, 0.01% Tween 20 (P1379, SigmaAldrich) and 2% donkey serum. Antibodies against TPRX1 1:200 (Merck, HPA044922) and H3.X/Y 1:200 (Active Motif, 61161) were diluted in blocking buffer and embryos incubated overnight at 4°C. They were then rinsed 3x in blocking buffer for at least 15 minutes per rinse, then incubated in secondary antibodies (Alexa Fluor conjugated, Molecular Probes) diluted 1:500 in blocking buffer for 1-2 hours at room temperature, rinsed in blocking buffer as previously, including DAPI to mark nuclei. After progression through increasing concentrations of Vectashield (H-1200, Vector Labs) in blocking buffer, they were mounted on glass slides in small drops of concentrated Vectashield. Coverslips were sealed with nail varnish. Multiple single optical sections were captured with a Carl Zeiss LSM780 microscope (63x oil-immersion objective).

### Immunofluorescence staining and imaging of human ESCs

Cells were plated on coverslips or cytospun onto poly-L-lysine coated glass slides (300rpm, 3min), washed with 1x PBS and fixed with 2% PFA in PBS for 30min at room temperature (RT). They were then permeabilized with 0.5% Triton X-100 in PBS for 1 hour at RT, blocked with 1% BSA, 0.05% Tween20 in PBS for 1 hour at RT, and incubated with primary antibody diluted in blocking solution for 1 hour at RT or o/n at 4°C. After washing in blocking solution for 30 min at RT, secondary antibodies were added for 1h at room temperature and cells were washed again. DNA was counterstained with DAPI (5mg/ mL in PBS) and slides were mounted using SlowFade Gold Antifade Mountant (Sigma Aldrich, S36937). The following antibodies and dilutions were used: TPRX1 1:200 (Merck, HPA044922), SOX2 1:300 (R&D systems, AF2018), H3.X/Y 1:200 (Active Motif, 61161). Secondary antibodies were Alexa Fluor conjugated and diluted 1:1000 (Molecular Probes). Single optical sections were captured with a Nikon A1-R (20x objective, 60x oil-immersion objective) or Carl Zeiss LSM780 microscope (63x oil-immersion objective). For visualization the images were pseudo-coloured and corrected for brightness and contrast (within the recommendations for scientific data) using Fiji (ImageJ 2.1.0/1.53c). Fluorescence semi-quantification analysis was performed with Volocity 6.3 (Improvision) on mid optical sections with manual segmentation.

### Quantitative Reverse Transcription PCR (qRT-PCR)

RNA from cells was isolated using RNeasy Mini kit (Qiagen, 74104) and treated with DNase (TURBO™ DNase 2U/ μL, Thermo Fisher Scientific, AM2238) according to manufacturer’s protocols. 1ug of DNAase-treated RNA was used for cDNA synthesis using the RevertAid First-Strand cDNA Synthesis Kit (Thermo Fisher Scientific, K1622). The cDNA was diluted (1:10-20) and used for qRT-PCR in technical triplicate using Brilliant III SYBR master mix (Agilent Technologies, 600882) and CFX384 Touch Real-Time PCR Detection Systems (BioRad). Relative levels of transcript expression were quantified by the comparative ΔΔCt method with normalisation to RPL19 levels. Primer sequences are available in Table S3.

### 10X Single-cell RNA-seq library preparation

For 10X single-cell RNA-seq, the cells were dissociated by incubating with 0.25% trypsin for 10min at 37°C, followed by trituration by pipetting with 200ul tip. The cells were resuspended in DMEM/F12 supplemented with 0.1% BSA, washed twice and then filtered through 30μm mesh. 16,000 cells were resuspended in 47μl DMEM/F12 supplemented with 0.04% BSA. Single-cell RNA-seq libraries were prepared in the Cancer Research UK Cambridge Institute Genomics Core Facility using the following: Chromium Single Cell 3’ Library & Gel Bead Kit v3 (10x Genomics, PN-1000075), Chromium Chip B Kit (10x Genomics, PN-1000073) and Chromium Single Cell 3’ Reagent Kits v3 User Guide (Manual Part CG000183 Rev C, 10X Genomics). Cell suspensions were loaded on the Chromium instrument with the expectation of collecting gel-beads emulsions containing single cells. RNA from the barcoded cells for each sample was subsequently reverse-transcribed in a C1000 Touch Thermal cycler (Bio-Rad) and all subsequent steps to generate single-cell libraries were performed according to the manufacturer’s protocol with no modifications. cDNA quality and quantity were measured with Agilent TapeStation 4200 (High Sensitivity 5000 ScreenTape) after which 25% of material was used for gene expression library preparation. Library quality was confirmed with Agilent TapeStation 4200 (High Sensitivity D1000 ScreenTape to evaluate library sizes) and Qubit 4.0 Fluorometer (ThermoFisher Qubit™ dsDNA HS Assay Kit to evaluate dsDNA quantity). Each sample was normalized and pooled in equal molar concentration. To confirm concentration, pool was qPCRed using KAPA Library Quantification Kit on QuantStudio 6 Flex before sequencing. Pool was sequenced on S2 flowcell on Illumina NovaSeq6000 sequencer with following parameters: 28 bp, read 1; 8 bp, i7 index; and 91 bp, read 2.

### Single-cell RNA-seq data analysis

Single-cell 10X RNA-seq samples were processed using the Cellranger count pipeline (v3.1.0) as Single Cell 3’ (v3) data using default parameters and raw sequencing reads were aligned to the GRCh38 human genome. The resulting data was filtered based on the distribution of the counts of RNA features, expression of mitochondrial genes, and percent largest genes. In our dataset, only cells with more than 2.500 and less than 10.500 genes, as well as less than 10% mitochondrial reads, and less than 10% largest gene percentage were kept. The data were normalised, scaled and variable features were identified using FindVariableFeatures function in Seurat (v4). PCA was performed based on variable features, UMAP was performed using the first 30 principal components and cells were clustered using Louvain at resolution 0.5. Differentially expressed genes between clusters were identified using the FindMarkers function (test.use=“roc”, only.pos = TRUE, logfc.threshold = 0.25). Plots were generated using DimPlot, DotPlot, VlnPlot, FeaturePlot, FeatureScatter, DoHeatmap and DimHeatmap functions. Cell cycle analysis was performed using the CellCycleScoring function with default parameters. The naïve hESCs dataset was integrated with human embryo data (E3 – E7, E-MTAB-3929) using the functions FindIntegrationAnchors and IntegrateData function. Merged data were normalized, scaled, and clustered.

### Transposable element analysis

Repeat family or subfamily ‘repeatomes’ were constructed using genome-wide annotation files of repeats generated by RepeatMasker (downloaded for the GRCh37 genome from the UCSC table browser). Individual instances of repeats were extracted from the genome sequence and concatenated together, whereby individual sequences were padded by NNNNN. To obtain FastQ files for single cells belonging to various clusters of interest, we used the cell level barcode files produced by Seurat (v4) as annotation files for subset10xbam (https://github.com/s-andrews/subset10xbam). This process uses the possorted BAM file outputted by CellRanger as well as the cell barcode annotations as input, and produces a new BAM file with entries only belonging to cells given in the annotation file as new output; during this process, the cell barcodes were added to the readID (option -- add_barcode). These files were then converted to FastQ format (using samtools fastq, version 1.11), and split into individual single-cell FastQ files using a custom script (https://github.com/FelixKrueger/scRepeats). These single-cell FastQ files were then aligned to various repeat genome sequences using Bowtie2 (default parameters), whereby any alignment to a repeat family was scored. The results were then converted to percentage of total reads that aligned to a given repeat class/family.

### Motif analysis

TF binding motifs overrepresented in our gene set were identified using RcisTarget with default parameters based on the hg19 database (version 1.10.0) (Aibar et al., 2017). Regions of 10kb centred around TSS of 8CLCs genes [top 200] were analysed and enriched motifs retrieved.

### RNA velocity analysis

RNA velocity analysis was based on initial processing of the 10x RNA sequencing data using velocyto (v0.17) (La Manno et al., 2018) and further analysis using scVelo (v0.2.4) (Bergen et al., 2020). We used velocyto to generate a loom file from 10x cellranger output data (‘velocyto run10x’, see also https://velocyto.org/velocyto.py/tutorial/cli.html) that differentiates between spliced, unspliced and ambiguous gene counts. This loom file was used for pre-processing and clustering via Seurat, and was then analysed using scVelo (via an h5ad output file, see also https://scvelo.readthedocs.io/). Scvelo (mode= ’stochastic’) was used to estimate trajectories based on spliced vs unspliced RNA for each cell and gene.

### scATAC-seq data analysis

10x multiome sequencing derived scATAC-seq data was analyzed using ArchR (v1.0.1) (Granja et al., 2021). Fragment files were loaded to generate arrow files, filtered (min TSS enrichment = 8, min fragments = 3000, max fragments = 1e7), integrated with cell identities from scRNA-seq and bigwig files from pseudobulks (getGroupBW) were exported for visualization using IGV.

## Data Availability

The single-cell RNA-seq data generated in this paper are deposited in Gene Expression Omnibus under the accession code GSE178379.

## Author contributions

J.T.S. conceived the project, designed and performed experiments, analysed the data and wrote the manuscript, with input from all authors. M.R performed single-cell RNA sequencing experiments, and provided expertise and helpful discussion. F.S. performed immunofluorescence experiments and analysis, and provided helpful discussions. S.L. helped with RNA velocity analysis. R.A. performed analysis of the 10x multiome data set. F.K. performed repeat alignment and analysis. J.N. provided embryo samples and performed immunofluorescence experiments. W.R. supervised the study and provided expertise and feedback.

## Acknowledgements

We thank all members of the W.R. laboratory for helpful discussions, and the Peter Rugg-Gunn laboratory for sharing reagents and stem cell expertise. We would also like to thank Christel Krueger and Simon Andrews for bioinformatics support, Simon Walker for imaging support and the members of the flow cytometry and the sequencing facility at the Babraham Institute for assistance. We would like to thank Katarzyna Kania and the CRUK sequencing facility for single-cell sequencing support. We thank Austin Smith for providing stem cell lines. We would like to acknowledge and thank the staff and patients at the assisted conception units at BMI Chelsfield Park and Leeds Fertility Hospitals. J.T.S. is supported by an EMBO Fellowship (ALTF 355-2019) and a Charles and Katharine Darwin Research Fellowship at Darwin College. S.L. was supported by the Donald Higham Fund at Peterhouse College. J.N. is supported by the University of Cambridge and BBSRC (BB/T007044/1). Research in W.R.’s lab is supported by the BBSRC (BBS/E/B/000C0422) and Wellcome Trust – Investigator award (210754/Z/18/Z). W.R. is a consultant and shareholder of Cambridge Epigenetix.

## References

Aibar, S., Gonzalez-Blas, C.B., Moerman, T., Huynh-Thu, V.A., Imrichova, H., Hulselmans, G., Rambow, F., Marine, J.C., Geurts, P., Aerts, J., et al. (2017). SCENIC: single-cell regulatory network inference and clustering. Nat Methods 14, 1083–1086.

Alda-Catalinas, C., Bredikhin, D., Hernando-Herraez, I., Santos, F., Kubinyecz, O., Eckersley-Maslin, M.A., Stegle, O., and Reik, W. (2020). A Single-Cell Transcriptomics CRISPR-Activation Screen Identifies Epigenetic Regulators of the Zygotic Genome Activation Program. Cell Syst 11, 25–41 e29.

Aoki, F., Worrad, D.M., and Schultz, R.M. (1997). Regulation of transcriptional activity during the first and second cell cycles in the preimplantation mouse embryo. Dev Biol 181, 296–307.

Bentsen, M., Goymann, P., Schultheis, H., Klee, K., Petrova, A., Wiegandt, R., Fust, A., Preussner, J., Kuenne, C., Braun, T., et al. (2020). ATAC-seq footprinting unravels kinetics of transcription factor binding during zygotic genome activation. Nat Commun 11, 4267.

Bergen, V., Lange, M., Peidli, S., Wolf, F.A., and Theis, F.J. (2020). Generalizing RNA velocity to transient cell states through dynamical modeling. Nat Biotechnol 38, 1408–1414.

Blakeley, P., Fogarty, N.M., del Valle, I., Wamaitha, S.E., Hu, T.X., Elder, K., Snell, P., Christie, L., Robson, P., and Niakan, K.K. (2015). Defining the three cell lineages of the human blastocyst by single-cell RNA-seq. Development 142, 3151–3165.

Braude, P., Bolton, V., and Moore, S. (1988). Human gene expression first occurs between the four- and eight-cell stages of preimplantation development. Nature 332, 459–461.

Bredenkamp, N., Yang, J., Clarke, J., Stirparo, G.G., von Meyenn, F., Dietmann, S., Baker, D., Drummond, R., Ren, Y., Li, D., et al. (2019). Wnt Inhibition Facilitates RNA-Mediated Reprogramming of Human Somatic Cells to Naive Pluripotency. Stem Cell Reports 13, 1083–1098.

De Iaco, A., Planet, E., Coluccio, A., Verp, S., Duc, J., and Trono, D. (2017). DUX-family transcription factors regulate zygotic genome activation in placental mammals. Nat Genet 49, 941–945.

Eckersley-Maslin, M.A., Svensson, V., Krueger, C., Stubbs, T.M., Giehr, P., Krueger, F., Miragaia, R.J., Kyriakopoulos, C., Berrens, R.V., Milagre, I., et al. (2016). MERVL/Zscan4 Network Activation Results in Transient Genome-wide DNA Demethylation of mESCs. Cell Rep 17, 179–192.

Evans, M.J., and Kaufman, M.H. (1981). Establishment in culture of pluripotential cells from mouse embryos. Nature 292, 154–156.

Falco, G., Lee, S.L., Stanghellini, I., Bassey, U.C., Hamatani, T., and Ko, M.S. (2007). Zscan4: a novel gene expressed exclusively in late 2-cell embryos and embryonic stem cells. Dev Biol 307, 539–550.

Granja, J.M., Corces, M.R., Pierce, S.E., Bagdatli, S.T., Choudhry, H., Chang, H.Y., and Greenleaf, W.J. (2021). ArchR is a scalable software package for integrative single-cell chromatin accessibility analysis. Nat Genet 53, 403–411.

Grow, E.J., Flynn, R.A., Chavez, S.L., Bayless, N.L., Wossidlo, M., Wesche, D.J., Martin, L., Ware, C.B., Blish, C.A., Chang, H.Y., et al. (2015). Intrinsic retroviral reactivation in human preimplantation embryos and pluripotent cells. Nature 522, 221–225.

Guo, G., von Meyenn, F., Rostovskaya, M., Clarke, J., Dietmann, S., Baker, D., Sahakyan, A., Myers, S., Bertone, P., Reik, W., et al. (2017). Epigenetic resetting of human pluripotency. Development 144, 2748–2763.

Guo, G., von Meyenn, F., Santos, F., Chen, Y., Reik, W., Bertone, P., Smith, A., and Nichols, J. (2016). Naive Pluripotent Stem Cells Derived Directly from Isolated Cells of the Human Inner Cell Mass. Stem Cell Reports 6, 437–446.

Hendrickson, P.G., Dorais, J.A., Grow, E.J., Whiddon, J.L., Lim, J.W., Wike, C.L., Weaver, B.D., Pflueger, C., Emery, B.R., Wilcox, A.L., et al. (2017). Conserved roles of mouse DUX and human DUX4 in activating cleavage-stage genes and MERVL/HERVL retrotransposons. Nat Genet 49, 925–934.

Huang, K., Maruyama, T., and Fan, G. (2014). The naive state of human pluripotent stem cells: a synthesis of stem cell and preimplantation embryo transcriptome analyses. Cell Stem Cell 15, 410–415.

Huang, Y., Kim, J.K., Do, D.V., Lee, C., Penfold, C.A., Zylicz, J.J., Marioni, J.C., Hackett, J.A., and Surani, M.A. (2017). Stella modulates transcriptional and endogenous retrovirus programs during maternal-to-zygotic transition. Elife 6.

Ishiuchi, T., Enriquez-Gasca, R., Mizutani, E., Boskovic, A., Ziegler-Birling, C., Rodriguez-Terrones, D., Wakayama, T., Vaquerizas, J.M., and Torres-Padilla, M.E. (2015). Early embryonic-like cells are induced by downregulating replication-dependent chromatin assembly. Nat Struct Mol Biol 22, 662–671.

Jagannathan, S., Shadle, S.C., Resnick, R., Snider, L., Tawil, R.N., van der Maarel, S.M., Bradley, R.K., and Tapscott, S.J. (2016). Model systems of DUX4 expression recapitulate the transcriptional profile of FSHD cells. Hum Mol Genet 25, 4419–4431.

Jiang, S., Williams, K., Kong, X., Zeng, W., Nguyen, N.V., Ma, X., Tawil, R., Yokomori, K., and Mortazavi, A. (2020). Single-nucleus RNA-seq identifies divergent populations of FSHD2 myotube nuclei. PLoS Genet 16, e1008754.

Jouhilahti, E.M., Madissoon, E., Vesterlund, L., Tohonen, V., Krjutskov, K., Plaza Reyes, A., Petropoulos, S., Mansson, R., Linnarsson, S., Burglin, T., et al. (2016). The human PRD-like homeobox gene LEUTX has a central role in embryo genome activation. Development 143, 3459–3469.

Kigami, D., Minami, N., Takayama, H., and Imai, H. (2003). MuERV-L is one of the earliest transcribed genes in mouse one-cell embryos. Biol Reprod 68, 651–654.

La Manno, G., Soldatov, R., Zeisel, A., Braun, E., Hochgerner, H., Petukhov, V., Lidschreiber, K., Kastriti, M.E., Lonnerberg, P., Furlan, A., et al. (2018). RNA velocity of single cells. Nature 560, 494–498.

Latham, K.E., and Schultz, R.M. (2001). Embryonic genome activation. Front Biosci 6, D748–759.

Lee, M.T., Bonneau, A.R., and Giraldez, A.J. (2014). Zygotic genome activation during the maternal-to-zygotic transition. Annu Rev Cell Dev Biol 30, 581–613.

Liu, L., Leng, L., Liu, C., Lu, C., Yuan, Y., Wu, L., Gong, F., Zhang, S., Wei, X., Wang, M., et al. (2019). An integrated chromatin accessibility and transcriptome landscape of human pre-implantation embryos. Nat Commun 10, 364.

Macfarlan, T.S., Gifford, W.D., Driscoll, S., Lettieri, K., Rowe, H.M., Bonanomi, D., Firth, A., Singer, O., Trono, D., and Pfaff, S.L. (2012). Embryonic stem cell potency fluctuates with endogenous retrovirus activity. Nature 487, 57–63.

Madissoon, E., Jouhilahti, E.M., Vesterlund, L., Tohonen, V., Krjutskov, K., Petropoulos, S., Einarsdottir, E., Linnarsson, S., Lanner, F., Mansson, R., et al. (2016). Characterization and target genes of nine human PRD-like homeobox domain genes expressed exclusively in early embryos. Sci Rep 6, 28995.

Maeso, I., Dunwell, T.L., Wyatt, C.D., Marletaz, F., Veto, B., Bernal, J.A., Quah, S., Irimia, M., and Holland, P.W. (2016). Evolutionary origin and functional divergence of totipotent cell homeobox genes in eutherian mammals. BMC Biol 14, 45.

Messmer, T., von Meyenn, F., Savino, A., Santos, F., Mohammed, H., Lun, A.T.L., Marioni, J.C., and Reik, W. (2019). Transcriptional Heterogeneity in Naive and Primed Human Pluripotent Stem Cells at Single-Cell Resolution. Cell Rep 26, 815–824 e814.

Nakamura, T., Okamoto, I., Sasaki, K., Yabuta, Y., Iwatani, C., Tsuchiya, H., Seita, Y., Nakamura, S., Yamamoto, T., and Saitou, M. (2016). A developmental coordinate of pluripotency among mice, monkeys and humans. Nature 537, 57–62.

Niakan, K.K., Han, J., Pedersen, R.A., Simon, C., and Pera, R.A. (2012). Human pre-implantation embryo development. Development 139, 829–841.

Peaston, A.E., Evsikov, A.V., Graber, J.H., de Vries, W.N., Holbrook, A.E., Solter, D., and Knowles, B.B. (2004). Retrotransposons regulate host genes in mouse oocytes and preimplantation embryos. Dev Cell 7, 597–606.

Percharde, M., Lin, C.J., Yin, Y., Guan, J., Peixoto, G.A., Bulut-Karslioglu, A., Biechele, S., Huang, B., Shen, X., and Ramalho-Santos, M. (2018). A LINE1-Nucleolin Partnership Regulates Early Development and ESC Identity. Cell 174, 391–405 e319.

Petropoulos, S., Edsgard, D., Reinius, B., Deng, Q., Panula, S.P., Codeluppi, S., Plaza Reyes, A., Linnarsson, S., Sandberg, R., and Lanner, F. (2016). Single-Cell RNA-Seq Reveals Lineage and X Chromosome Dynamics in Human Preimplantation Embryos. Cell 165, 1012–1026.

Pontis, J., Planet, E., Offner, S., Turelli, P., Duc, J., Coudray, A., Theunissen, T.W., Jaenisch, R., and Trono, D. (2019). Hominoid-Specific Transposable Elements and KZFPs Facilitate Human Embryonic Genome Activation and Control Transcription in Naive Human ESCs. Cell Stem Cell 24, 724–735 e725.

Resnick, R., Wong, C.J., Hamm, D.C., Bennett, S.R., Skene, P.J., Hake, S.B., Henikoff, S., van der Maarel, S.M., and Tapscott, S.J. (2019). DUX4-Induced Histone Variants H3.X and H3.Y Mark DUX4 Target Genes for Expression. Cell Rep 29, 1812–1820 e1815.

Rodriguez-Terrones, D., Gaume, X., Ishiuchi, T., Weiss, A., Kopp, A., Kruse, K., Penning, A., Vaquerizas, J.M., Brino, L., and Torres-Padilla, M.E. (2018). A molecular roadmap for the emergence of early-embryonic-like cells in culture. Nat Genet 50, 106–119.

Roode, M., Blair, K., Snell, P., Elder, K., Marchant, S., Smith, A., and Nichols, J. (2012). Human hypoblast formation is not dependent on FGF signalling. Dev Biol 361, 358–363.

Rostovskaya, M., Stirparo, G.G., and Smith, A. (2019). Capacitation of human naive pluripotent stem cells for multi-lineage differentiation. Development 146.

Shen, H., Yang, M., Li, S., Zhang, J., Peng, B., Wang, C., Chang, Z., Ong, J., and Du, P. (2021). Mouse totipotent stem cells captured and maintained through spliceosomal repression. Cell 184, 2843–2859 e2820.

Stirparo, G.G., Boroviak, T., Guo, G., Nichols, J., Smith, A., and Bertone, P. (2018). Integrated analysis of single-cell embryo data yields a unified transcriptome signature for the human pre-implantation epiblast. Development 145.

Sumi, T., Oki, S., Kitajima, K., and Meno, C. (2013). Epiblast ground state is controlled by canonical Wnt/beta-catenin signaling in the postimplantation mouse embryo and epiblast stem cells. PLoS One 8, e63378.

Svoboda, P., Stein, P., Anger, M., Bernstein, E., Hannon, G.J., and Schultz, R.M. (2004). RNAi and expression of retrotransposons MuERV-L and IAP in preimplantation mouse embryos. Dev Biol 269, 276–285.

Takashima, Y., Guo, G., Loos, R., Nichols, J., Ficz, G., Krueger, F., Oxley, D., Santos, F., Clarke, J., Mansfield, W., et al. (2014). Resetting transcription factor control circuitry toward ground-state pluripotency in human. Cell 158, 1254–1269.

Tarkowski, A.K. (1959). Experiments on the development of isolated blastomers of mouse eggs. Nature 184, 1286–1287.

Theunissen, T.W., Friedli, M., He, Y., Planet, E., O’Neil, R.C., Markoulaki, S., Pontis, J., Wang, H., Iouranova, A., Imbeault, M., et al. (2016). Molecular Criteria for Defining the Naive Human Pluripotent State. Cell Stem Cell 19, 502–515.

Theunissen, T.W., Powell, B.E., Wang, H., Mitalipova, M., Faddah, D.A., Reddy, J., Fan, Z.P., Maetzel, D., Ganz, K., Shi, L., et al. (2014). Systematic Identification of Culture Conditions for Induction and Maintenance of Naive Human Pluripotency. Cell Stem Cell 15, 524–526.

Thomson, J.A., Itskovitz-Eldor, J., Shapiro, S.S., Waknitz, M.A., Swiergiel, J.J., Marshall, V.S., and Jones, J.M. (1998). Embryonic stem cell lines derived from human blastocysts. Science 282, 1145–1147.

Vassena, R., Boue, S., Gonzalez-Roca, E., Aran, B., Auer, H., Veiga, A., and Izpisua Belmonte, J.C. (2011). Waves of early transcriptional activation and pluripotency program initiation during human preimplantation development. Development 138, 3699–3709.

Wang, J., Xie, G., Singh, M., Ghanbarian, A.T., Rasko, T., Szvetnik, A., Cai, H., Besser, D., Prigione, A., Fuchs, N.V., et al. (2014). Primate-specific endogenous retrovirus-driven transcription defines naive-like stem cells. Nature 516, 405–409.

Xue, Z., Huang, K., Cai, C., Cai, L., Jiang, C.Y., Feng, Y., Liu, Z., Zeng, Q., Cheng, L., Sun, Y.E., et al. (2013). Genetic programs in human and mouse early embryos revealed by single-cell RNA sequencing. Nature 500, 593–597.

Yan, L., Yang, M., Guo, H., Yang, L., Wu, J., Li, R., Liu, P., Lian, Y., Zheng, X., Yan, J., et al. (2013). Single-cell RNA-Seq profiling of human preimplantation embryos and embryonic stem cells. Nat Struct Mol Biol 20, 1131–1139.

Yao, Z., Snider, L., Balog, J., Lemmers, R.J., Van Der Maarel, S.M., Tawil, R., and Tapscott, S.J. (2014). DUX4-induced gene expression is the major molecular signature in FSHD skeletal muscle. Hum Mol Genet 23, 5342–5352.

Zalzman, M., Falco, G., Sharova, L.V., Nishiyama, A., Thomas, M., Lee, S.L., Stagg, C.A., Hoang, H.G., Yang, H.T., Indig, F.E., et al. (2010). Zscan4 regulates telomere elongation and genomic stability in ES cells. Nature 464, 858–863.

