## Supplementary Figures S1-S6 for "Modelling human zygotic genome activation in 8C-like cells *in vitro*"

This PDF file includes:

Supplemental Figures S1 – S6

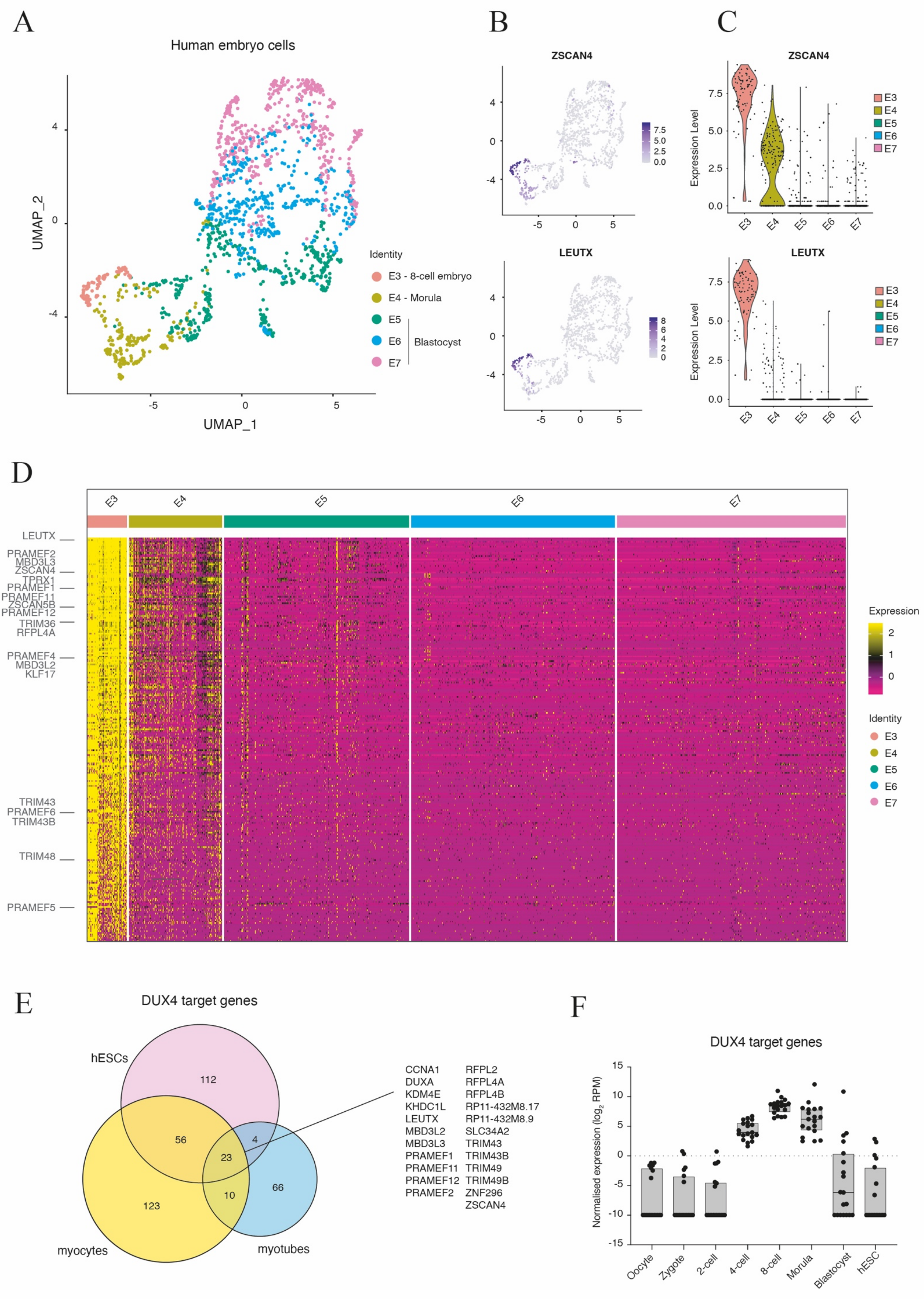


Supplemental Figure 1

**Figure S1.** (A) UMAP of human 8-cell (E3), morula (E4) and blastocyst (E5 – E7) stage embryo cells based on single cell RNA expression data (Petropoulos et al., 2016). (B) UMAP and (C) Violin plots of normalized, scaled *ZSCAN4* and *LEUTX* expression in human pre-implantation embryo cells. (D) Heatmap of expression of 8-cell embryo signature genes (rows) in human pre-implantation embryo cells (E3 – E7, columns). ZGA markers are highlighted. (E) Differentially expressed genes that are upregulated upon *DUX4* overexpression in primed hPSCs (magenta), myocytes (yellow), and endogenous *DUX4* upregulation in human patient myotubes (blue) (Hendrickson et al., 2017; Jiang et al., 2020; Yao et al., 2014). Shared genes across all three datasets are highlighted. (F) Normalized expression of *DUX4* target genes during human pre-implantation development (oocyte to blastocyst) and in hESCs (Yan et al., 2013).

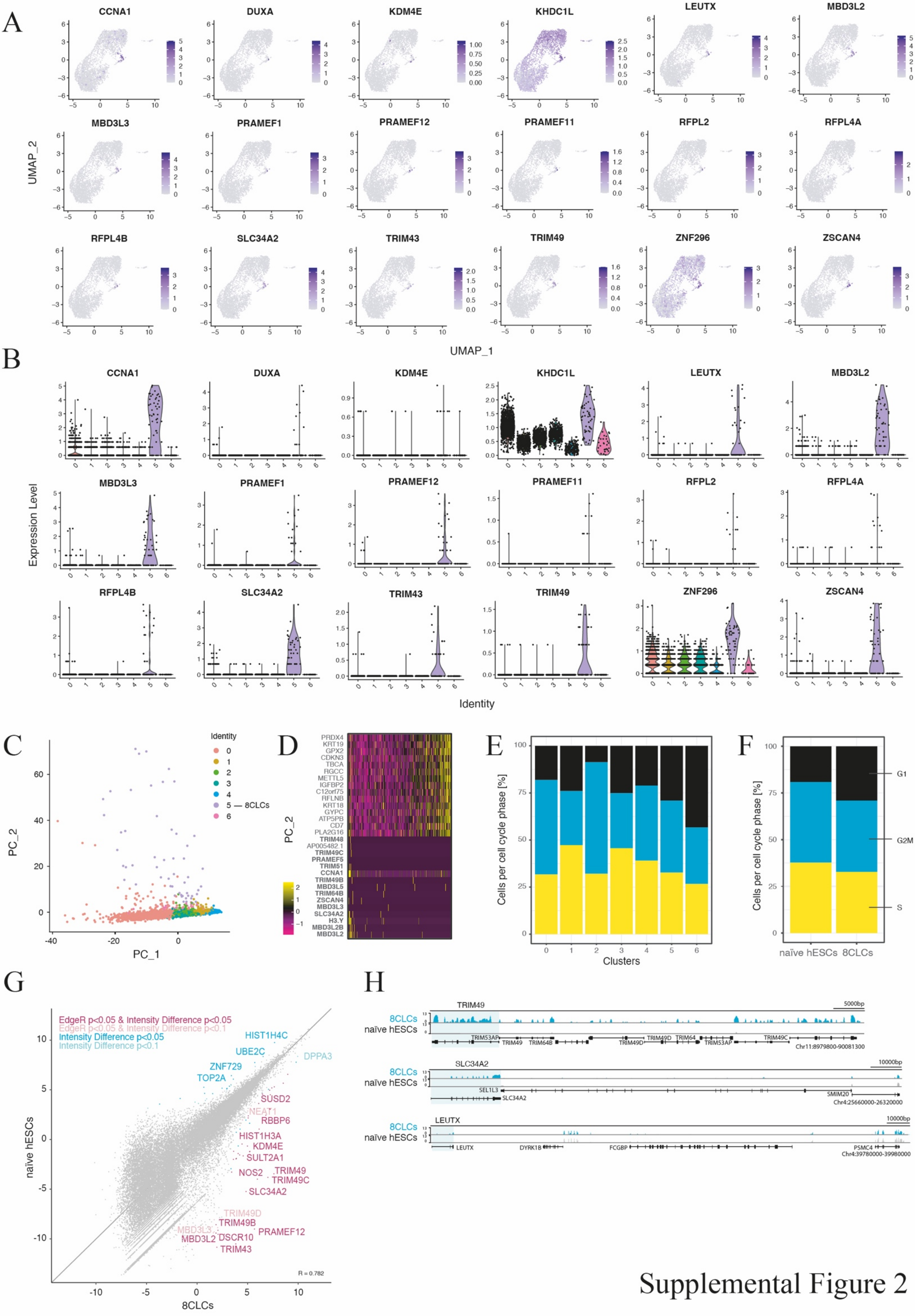


Supplemental Figure 2

**Figure S2.** (A) UMAP and (B) Violin plots of selected ZGA markers in clustered human naïve ESCs, based on normalized, scaled single cell RNA expression levels. Clusters were determined as in Figure 1 (B). (C) PC1 and PC2 of clustered naïve human HNES1 ESCs. (D) PC 2 loadings of naïve HNES1 are shown. ZGA marker genes in PC2 are highlighted in bold. (E) Cell cycle analysis of single cell transcriptome data in naïve hESCs clusters (cluster 0 – 6) or (F) specifically 8CLCs as compared to naïve hESCs. (G) Differentially expressed genes, as defined by EdgeR or Intensity Difference analysis, between 8CLCs (n=5/480, based on the expression of ZGA markers) and naïve H9 PSCs cultured in t2iLGö (Messmer et al., 2019). (H) Genome browser tracks of pseudo-bulked ZGA marker gene expression in 8CLCs (as in G) and naïve H9 hPSCs (Messmer et al., 2019).

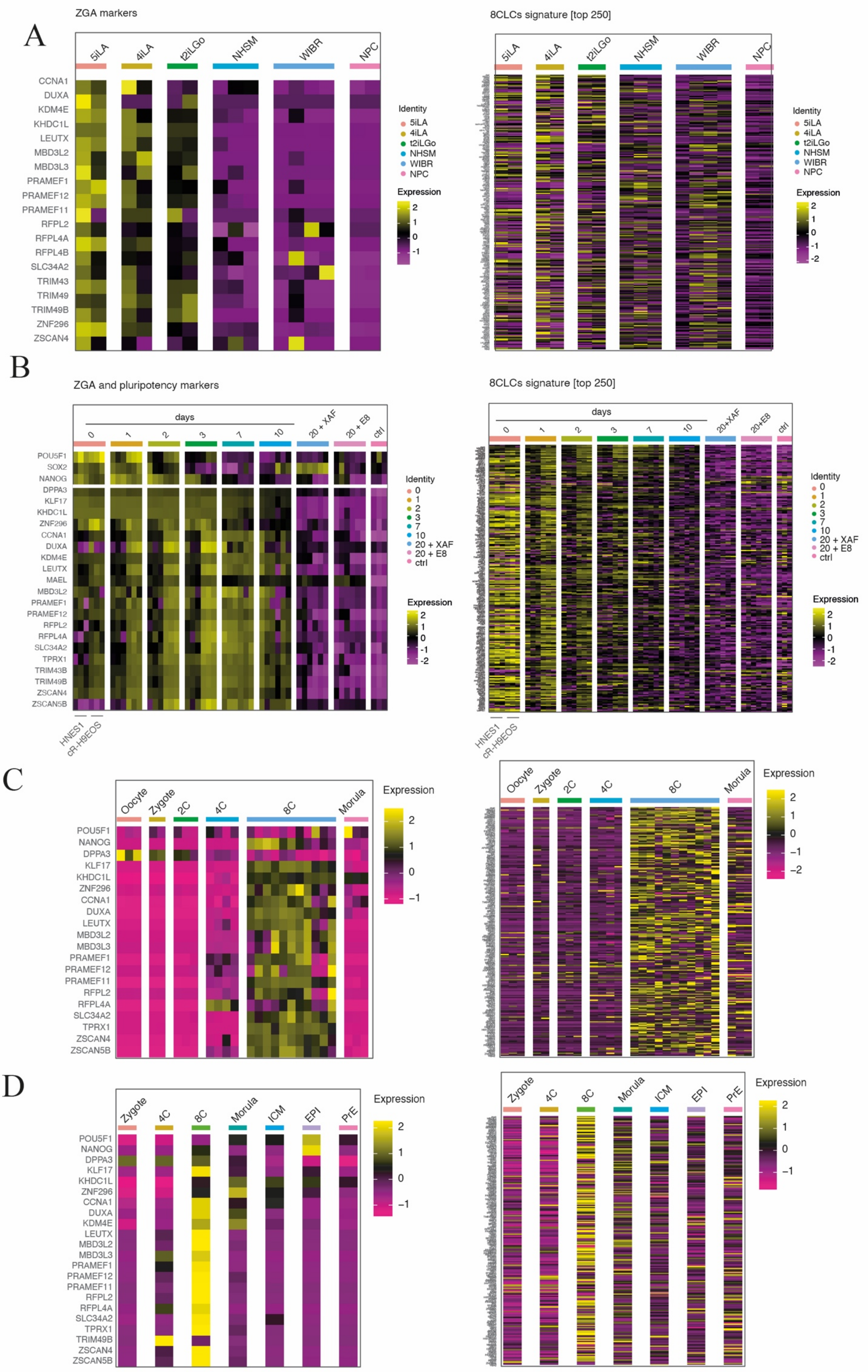


Supplemental Figure 3

**Figure S3.** (A) Heatmap of ZGA markers (left panel) and 8CLCs signature genes (right panel) of naïve (5iLA, 4iLA, t2iLGö), pseudo-naïve (NHSM), primed (WIBR) and differentiated cells (NPCs, neuronal progenitor cells). The analysis was done based on bulk RNA expression data. (B) Heatmap of pluripotency genes (i.e. *POU5F1, SOX2, NANOG*), ZGA markers (left panel), and 8CLCs signature genes (right panel) in two independent naïve PSC lines (HNES1, cR-H9EOS in triplicate each). Cells were transitioned from the naïve into the primed state, in two different primed media (E8, XAF) (Rostovskaya et al., 2019). The days indicate the days of transition (day 0 – day 10, d20). Gene expression of control primed H9 cells is shown as well (ctrl). (C), (D) Heatmap of pluripotency genes (*POU5F1, NANOG*), ZGA markers (left panels), and 8CLCs signature genes (right panels) in human pre-implantation embryos from two different studies (C) (Xue et al., 2013) (D) (Stirparo et al., 2018).

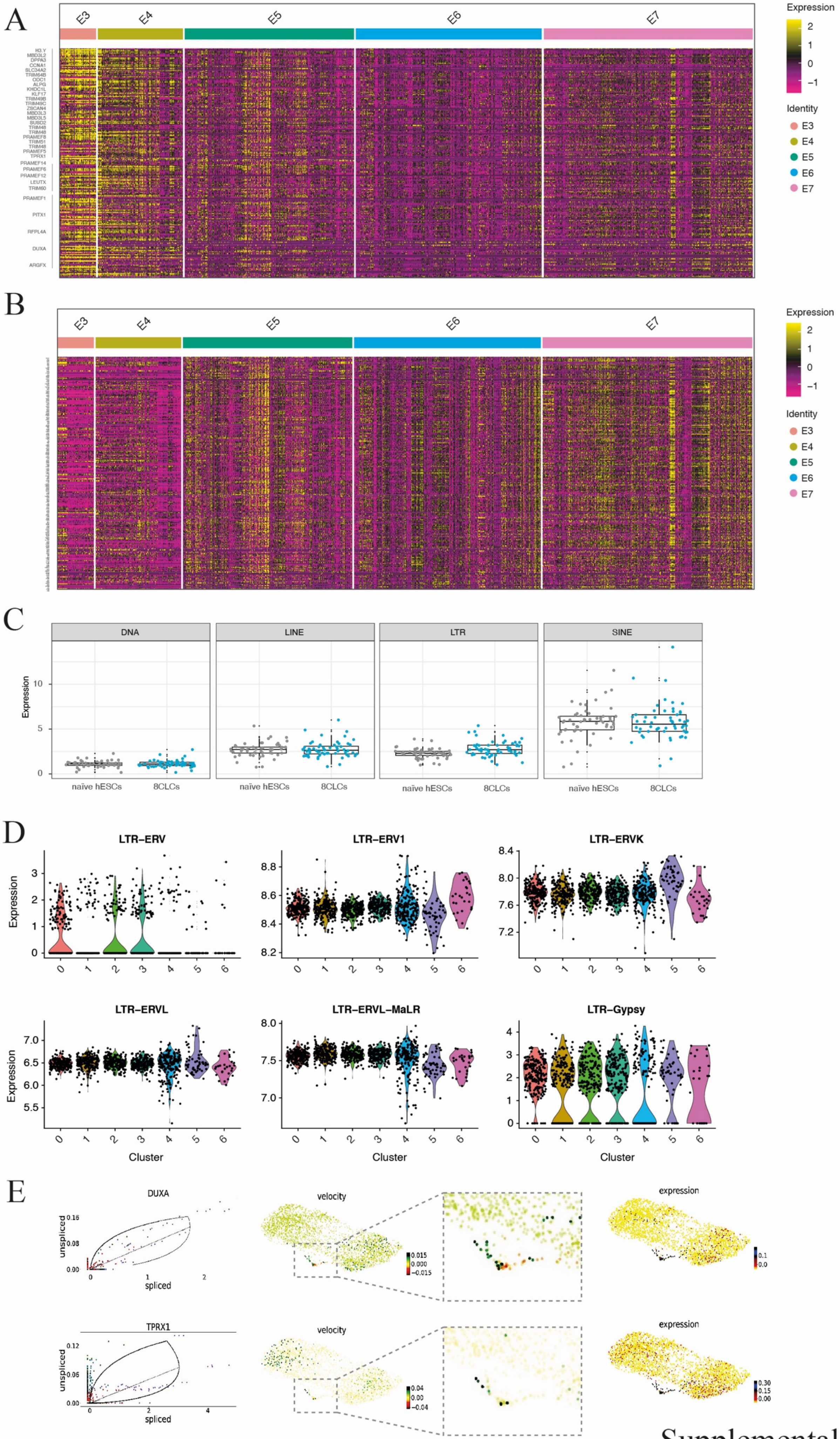


Supplemental Figure 4

**Figure S4.** (A) 8CLCs signature and (B) naïve hPSCs marker gene expression that distinguishes them from 8CLCs, in human embryos (E3 – E7) (Petropoulos et al., 2016). (C) DNA, LTR, LINE, and SINE transposable element expression in naïve hESCs and 8CLCs. Reads are depicted as percentage of total reads. (D) Normalized reads of LTR family transposons expressed in 8CLCs and naïve hESCs from single cell RNA sequencing data (for clusters see Figure 1B). (E) Left panel: RNA velocity analysis of *DUXA* and *TPRX1* in individual 8CLCs and naïve hESCs; steady-state ratios (black lines), overall dynamics (black curve) and ratios of unspliced vs. spliced mRNA in single cells are shown; middle panel: RNA velocity of marker genes; right panel: RNA expression levels.

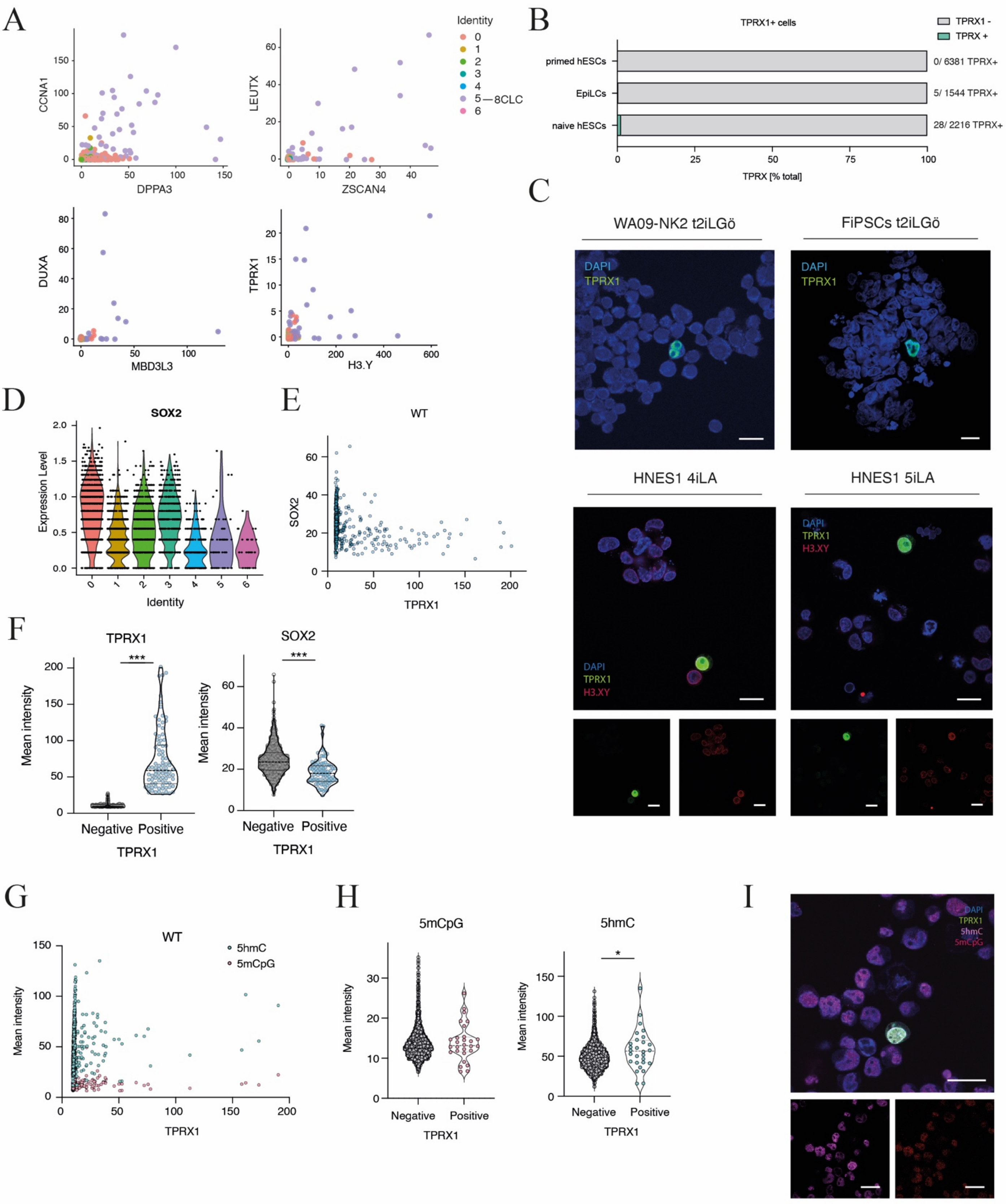


Supplemental Figure 5

**Figure S5.** (A) Scatter plots of 8CLC marker gene expression in naïve hESCs and 8CLC clusters. (B) Quantification of TPRX1+ naïve hESCs, EpiLCs and primed ESCs as detected by immunofluorescence staining. Percentages are shown in the graph and total numbers are written on the right-hand side. (C) Upper panel: IF staining of TPRX1 in reprogrammed naïve H9 PSCs (left) and fibroblast derived naïve iPSCs (FiPSCs, right) cultured in t2iLGö. Bottom panel: IF staining of TRPX1 and H3.X/Y in naïve HNES1 cells cultured in 4iLA (left) or 5iLA (right). Scale bar, 20µm. (D) Violin plot of normalized, scaled *SOX2* expression in clustered naïve human ESCs. (E) TPRX1 and SOX2 mean fluorescent signal intensity per cell (mid optical slice) in antibody stained, cytospun naïve HNES1 cells cultured in PXGL. TPRX1-positive cells and control cells were selected to assess SOX2 protein levels. (F) Mean TPRX1 (left) and SOX2 (right) intensity levels in TPRX1-positive and TPRX1-negative selected HNES1 cells. Images were taken from four independent experiments and pooled. p-value *P<0.05, **P<0.01, ***P<0.001, absence of stars (non-significant, ns): p-value>0.05; two tailed Mann-Whitney U test. (G) TRPX1 and 5hmC/ 5mCpG mean fluorescent signal intensity per cell (mid optical slice) in antibody stained, cytospun naïve HNES1 cells cultured in PXGL. (H) Mean 5mCpG (left) and 5hmC (right) intensity levels in TPRX1-positive and TPRX1-negative HNES1 cells. Images were taken from at least two experiments and pooled. p-value *P<0.05, **P<0.01, ***P<0.001, absence of stars (non-significant, ns): p-value>0.05; unpaired, two tailed Mann-Whitney U test. (I) IF staining of TRPX1, 5hmC and 5mCpG in naïve HNES1 cells cultured in PXGL. Scale bar, 20µm.

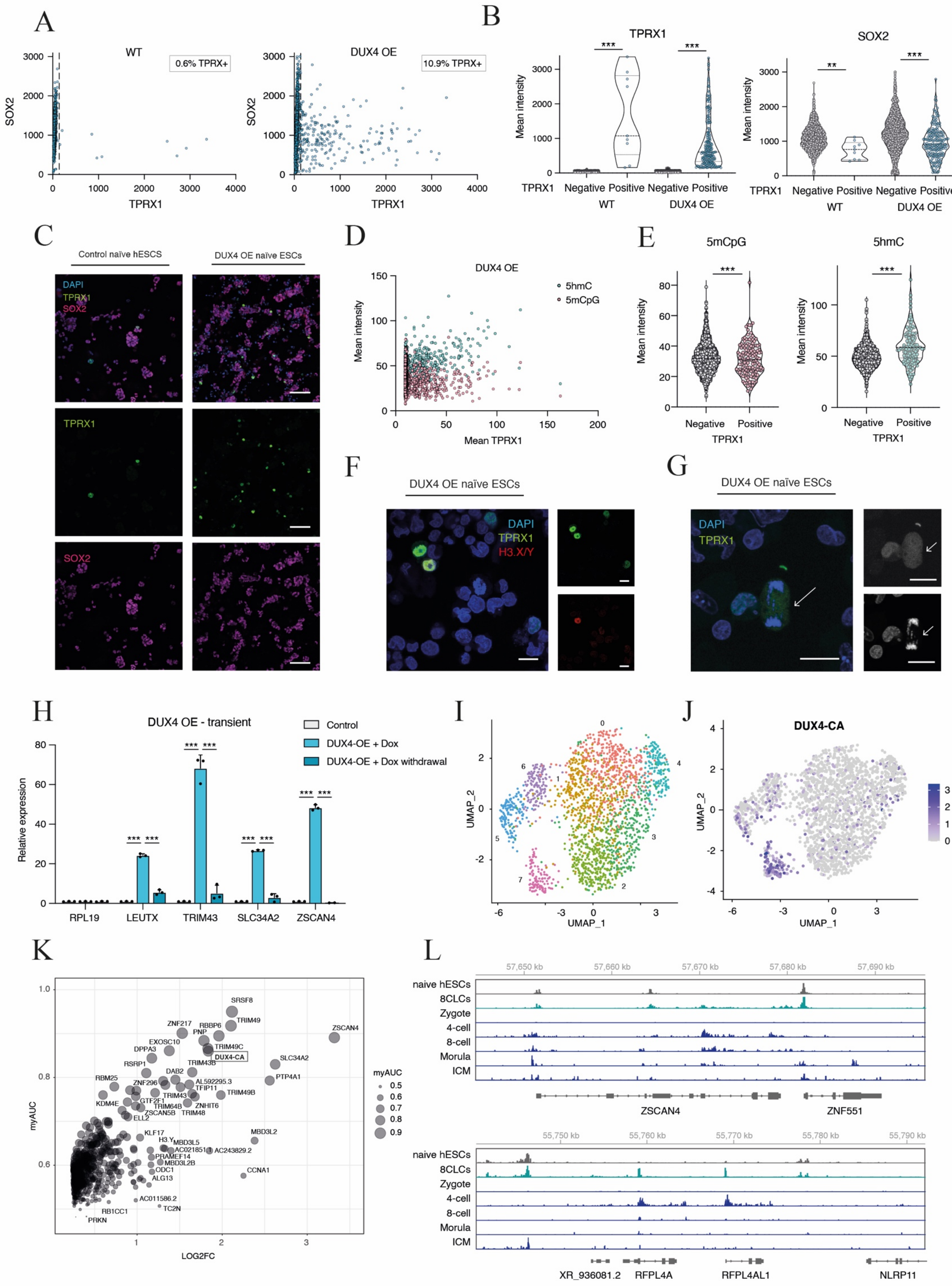


Supplemental Figure 6

**Figure S6.** (A) TPRX1 and SOX2 mean fluorescent signal intensity per cell in antibody-stained control (WT) and *DUX4* overexpressing (*DUX4* OE) naïve HNES1 cells cultured in PXGL and plated on coverslips. The percentage of TPRX1-positive cells is indicated on the top right of the graph, the cut-off is shown as dashed line (x=150). (B) Mean TPRX1 (left) and SOX2 (right) intensity levels in TPRX1-positive and TPRX1-negative control (WT) and *DUX4* overexpressing (*DUX4* OE) HNES1 cells; two independent experiments. Cut-off as in (A). p-value $*P<0.05$, $**P<0.01$, $***P<0.001$, absence of stars (non-significant, ns): p-value$>0.05$; two tailed Mann-Whitney U test. (C) IF staining of TPRX1 and SOX2 in control and *DUX4* overexpressing HNES1 cells plated on coverslips. Scale bar, 100µm. (D) TRPX1 and 5hmC/ 5mCpG mean fluorescent signal intensity per cell (mid optical slice) in antibody stained, cytospun *DUX4*-overexpression HNES1 cells cultured in PXGL. (E) Mean 5mCpG (left) and 5hmC (right) intensity levels in TPRX1-positive and TPRX1-negative *DUX4*-overexpressing HNES1 cells; two independent experiments. p-value $*P<0.05$, $**P<0.01$, $***P<0.001$, absence of stars (non-significant, ns): p-value$>0.05$; nonparametric Mann-Whitney U test. (F) IF staining of TPRX1 and H3.X/Y in *DUX4* overexpressing HNES1 cells cytospun onto coverslips. Scale bar, 20µm (G) IF staining of TPRX1 in *DUX4* overexpressing cytospun HNES1 cells. Dividing TPRX1-positive cells (anaphase chromosomes) are indicated (see arrows). Scale bar, 20µm. (H) Expression levels of ZGA marker genes measured by RT-qPCR in DUX4 overexpressing (DUX4-OE + Dox, 24h) cells as well as upon Dox withdrawal (DUX-OE + Dox withdrawal, 24h + 72h). Data are shown as mean ± SD (n = 3 technical replicates) of fold-change compared to control HNES1, and are representative of three independent experiments. p-value $*P<0.05$, $**P<0.01$, $***P<0.001$, absence of stars (non-significant, ns): p-value$>0.05$; unpaired, two tailed Student's t-test. (I) UMAP of clustered human naïve ESCs and 8CLCs (cluster 7). (J) Normalized scaled *DUX4-CA* expression in clustered hESCs and 8CLCs visualized on a UMAP. (K) Marker gene expression signature of 8CLCs (cluster 7 see S6I). AUC, area under curve; LOG2FC, log2 fold-change. (L) Genome browser views of accessibility at genomic loci of the ZGA markers *ZSCAN4* (data range 0.0 – 0.2 stem cells, 0.0 – 1.0 embryo data) and *RFPL4A* (data range 0.0 – 0.1 stem cells, 0.0 – 1.0 embryo data) in naïve hESCs (grey) and 8CLCs (petrol) and human embryos (blue) (Liu et al., 2019).
